# Decorrelation by gain control in the mouse olfactory bulb

**DOI:** 10.64898/2026.05.07.722633

**Authors:** Sina Tootoonian, Yikai Yang, Andreas T Schaefer, Tobias Ackels

## Abstract

Classifying sensory stimuli requires neural circuits to transform overlapping inputs into representations that are more distinct and decodable. Odour representations in the olfactory bulb (OB) have been reported to decorrelate from sensory input to output, a transformation thought to enhance odour separability. What has been missing is a systematic quantification of this transformation across odours, and a mechanistic account of how it can be implemented by OB circuitry. Using *in vivo* two-photon calcium imaging, we recorded glomerular input and output responses to a chemically diverse panel of 47 odours in mice. We observed a robust decorrelation of odour-evoked activity patterns from input to output: correlations between odour representations were consistently reduced, while overall response variance was preserved. This transformation increased the dimensionality of the population code and improved the separability of odour representations. Consistent with this, classification analyses showed that odour identity could be decoded with significantly higher accuracy at the output. To understand the mechanisms underlying this transformation, we developed a linear model of the OB with different levels of connectivity. Models in which each channel underwent only self-gain modulation reproduced the observed decorrelation nearly as well as models with fully unconstrained lateral connectivity. Analysis revealed that channels contributing unique information were selectively amplified, while redundant channels were attenuated. Together, our results show that the OB implements a measurable input–output transformation that enhances odour separability. By combining systematic recordings with computational modelling, we found that feedforward gain modulation serves as a simple and scalable mechanism capable of explaining the observed decorrelation, reframing how we understand the functional architecture of early olfactory processing.

## Introduction

Every organism inhabits a sensory world rich in information that must be efficiently distinguished and acted upon (Dangles et al., 2009). Neural systems face the challenge of transforming complex and often overlapping inputs into representations that are separable enough to guide behaviour. Studying how this transformation unfolds across processing stages provides insight into the general principles by which the nervous system extracts and organizes information.

A key feature of sensory processing is the correlation structure of neural activity, which reflects how sensory inputs are transformed into representations that can be effectively distinguished by downstream circuits. In order to support behaviourally relevant discrimination, the correlation structure is thought to be shaped to emphasize features of the natural environment that matter to the organism (Li & Liberles, 2015). However, excessive similarity between population responses can limit the discriminability of sensory inputs (Bishop, 1995; Moreno-Bote et al., 2014). In the mammalian olfactory bulb (OB), olfactory sensory neurons (OSNs) converge onto glomeruli to form a spatial map of receptor identity (Vassar et al., 1994). Glomerular output is relayed by mitral and tufted (M/T) cells, which interact with a dense network of inhibitory interneurons (Burton, 2017; Fukunaga et al., 2014; Mori et al., 1999; Parrish-Aungst et al., 2007; Shepherd, 2004). These circuits transform sensory input before transmission to higher brain regions (Arneodo et al., 2018; Chae et al., 2019; Igarashi et al., 2012; Nagayama, 2010; Storace & Cohen, 2017).

Decorrelation of odour representations has been observed across species including flies (Bhandawat et al., 2007; Lin et al., 2014), zebrafish (Friedrich & Laurent, 2001, 2004; Friedrich & Wiechert, 2014; Wanner & Friedrich, 2020), and mice (Gschwend et al., 2015; Otazu et al., 2015). Inhibitory circuits have been proposed as a mechanism for such decorrelation, and in larval zebrafish, disynaptic feedforward inhibition has been shown to be both sufficient and necessary to decorrelate odour representations (Wanner & Friedrich, 2020). In mice, interneurons contribute to decorrelation in mixture processing and discrimination behaviour (Abraham et al., 2010; Gschwend et al., 2015; Lepousez et al., 2014; Shimshek et al., 2005), and top-down cortical feedback can also reduce correlations among OB output patterns (Otazu et al., 2015). In addition to spatial mechanisms, temporal dynamics further shape decorrelation, with odour responses becoming progressively less correlated on timescales of hundreds of milliseconds (Friedrich & Laurent, 2001, 2004; Gschwend et al., 2015; Wanner & Friedrich, 2020).

While these studies demonstrate that odour representations decorrelate from input to output, two central questions remain. First, how large and systematic is this transformation across a broad odour space? Second, what circuit motifs are necessary and sufficient to generate the observed decorrelation in mice?

To address these questions, we combined experimental and computational approaches. Using *in vivo* two-photon calcium imaging, we measured glomerular responses to a large and chemically diverse odour panel at both the input and output levels of the OB. We found robust decorrelation, increased dimensionality, and enhanced decoding accuracy at the output stage. To uncover the mechanisms underlying these transformations, we developed a linear model of the OB consisting of glomerular inputs, mitral cells, and inhibitory connectivity between them. Remarkably, channel-specific gain modulation alone reproduced the observed decorrelation nearly as well as unconstrained connectivity, implying that feedforward scaling rather than lateral interactions underlies much of the OB’s input–output transformation.

Thus, our experimental and modelling results offer a concrete mechanistic account of how the OB enhances odour separability.

## Results

### Measuring glomerular odour response profiles from input and output activity

Axons of olfactory sensory neurons (OSNs) that express the same receptor converge onto glomeruli located in the outer layer of the olfactory bulb (OB). These glomeruli serve as the initial site of synaptic input to projection neurons, called mitral and tufted cells (M/T cells), which receive excitatory input through their apical dendrites. Recording neural activity in the glomerular layer therefore allows measurement of a summed population signal. For the sensory input, this corresponds to OSNs expressing the same receptor; for outputs, it reflects the summed activity of M/T cells associated with the same glomerulus.

To measure glomerular odour response profiles, we recorded input (n = 5) and output (n = 12) activity from the dorsal OB in separate head-fixed mice using *in vivo* two-photon calcium imaging (**Figure 1A, left**). Mice were presented with a panel of 47 monomolecular odours spanning a broad chemical range and mineral oil as a blank stimulus (**Figure 1B, S1.1, S1.2**) (Burton et al., 2022; Chae et al., 2019; Pashkovski et al., 2020). We used triple-transgenic mouse lines expressing GCaMP6f to capture calcium dynamics of either the sensory input in axons of sensory neurons (crossed with an OMP-Cre driver line, **Figure 1C, left**) or output in apical tufts of M/T cells (crossed with a Tbet-Cre driver line, **Figure 1C, right**), alongside a fluorescently tagged M72 or O174 glomerulus. The genetically targeted glomerulus served as an anatomical landmark to facilitate imaging from a similar area of the OB across experiments (**Figure 1A, right**). Glomerular regions of interest (ROIs) were manually selected from maximum-intensity projections for the imaged fields of view (OSN axons for input; **Figure 1D**, M/T cell dendrites for output; **Figure 1E**). Calcium signals evoked by odour stimulation were then extracted for further analysis (see Methods).

**Figure 1:**
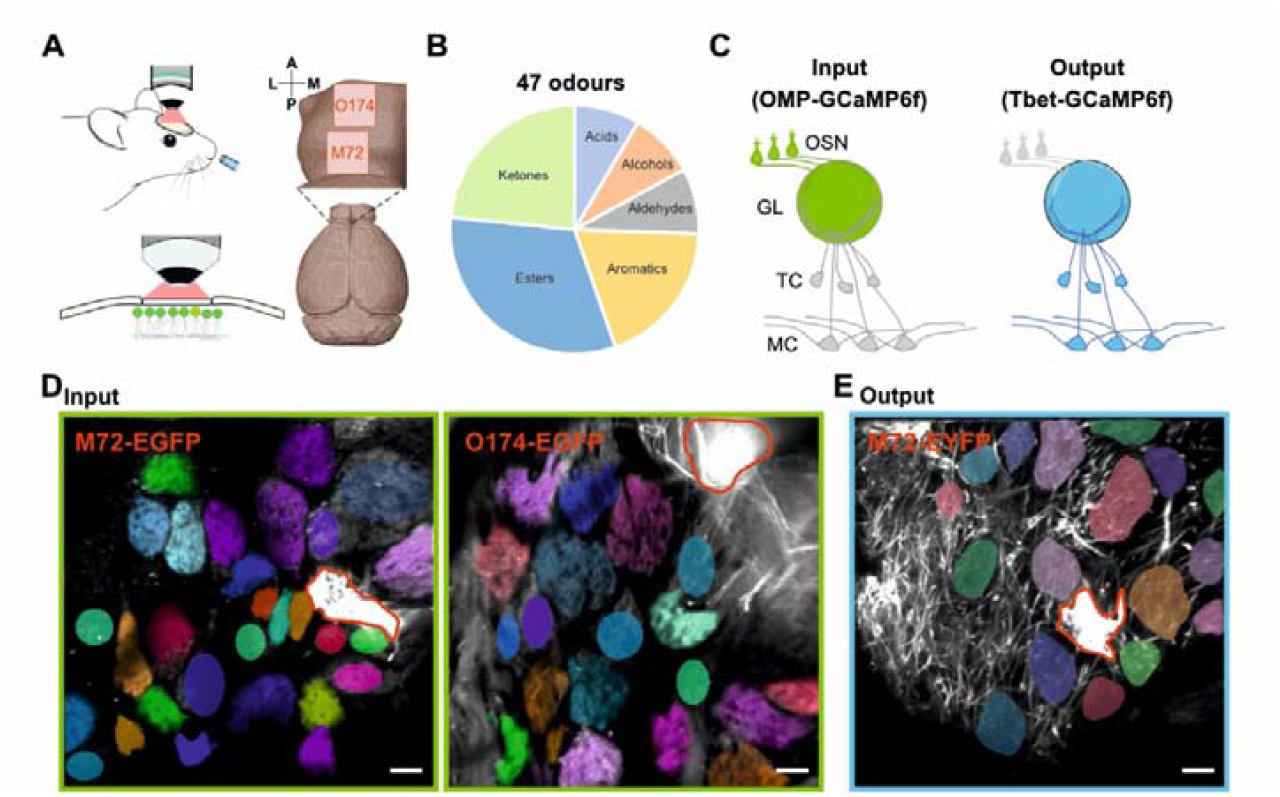
Measuring glomerular odour response profiles from input and output activity. **(A)** *In vivo* two-photon imaging setup for olfactory bulb glomerular calcium imaging in head-fixed mice. Imaging areas are located around a genetically targeted glomerulus. **(B)** Odour panel consisting of 47 monomolecular odours spanning six chemical classes. **(C)** Schematic representation of the glomerular calcium signal source using different transgenic animals. Calcium signals were recorded from the sensory input (green; OMP-GCaMP6f) and output (blue; Tbet-GCaMP6f) in separate animals. **(D)** GCaMP6f fluorescence from olfactory sensory neuron axons (8000 frames maximum projections) recorded in the glomerular layer of OMP-GCaMP6f mice. EGFP expression in sensory axons allows identification of genetically labelled glomeruli (red contour, O174; n = 3 or M72; n = 2 animals; coloured areas represent manually selected glomeruli). **(E)** Same as (D) but the GCaMP6f signal in the glomerular layer originates from output neuron dendrites (n = 12 animals). Scale bars, 50 μm. OSN: olfactory sensory neuron, GL: glomerulus, TC: tufted cell, MC: mitral cell.

### Input and output responses reveal structured and diverse odour representations

Glomerular response profiles are determined by the molecular receptive range of individual olfactory receptors. Here, we measured the odour-evoked activity in olfactory bulb glomeruli as a summed signal. Using a panel of 47 chemically diverse monomolecular odours (**Figure S1.1**; (Burton et al., 2022; Chae et al., 2019; Pashkovski et al., 2020)), we generated response profiles for 148 individual glomeruli for the sensory input (**Figure 2A-C**) and 167 glomeruli for the output (**Figure 2D-F**). Odour responses at both input and output levels varied systematically with odour identity and exhibited diverse patterns across glomeruli (**Figure 2**). Individual glomeruli responded consistently over repeated presentations of the same odour (**Figure 2A,D**; Odour 9, 4-phenyl-2-butanone). To visualise population-level response dynamics, we plotted time series of average responses for all recorded glomeruli as heatmaps (**Figure 2B,E**), revealing a wide range of response kinetics. Static response maps summarising activity across all glomeruli and odours were derived from z-scored fluorescence changes integrated over a 5-second window (**Figure 2C,F**). The odour order was determined using hierarchical clustering based on the Euclidean distance of the input data and was kept consistent across input and output datasets (see Methods).

**Figure 2:**
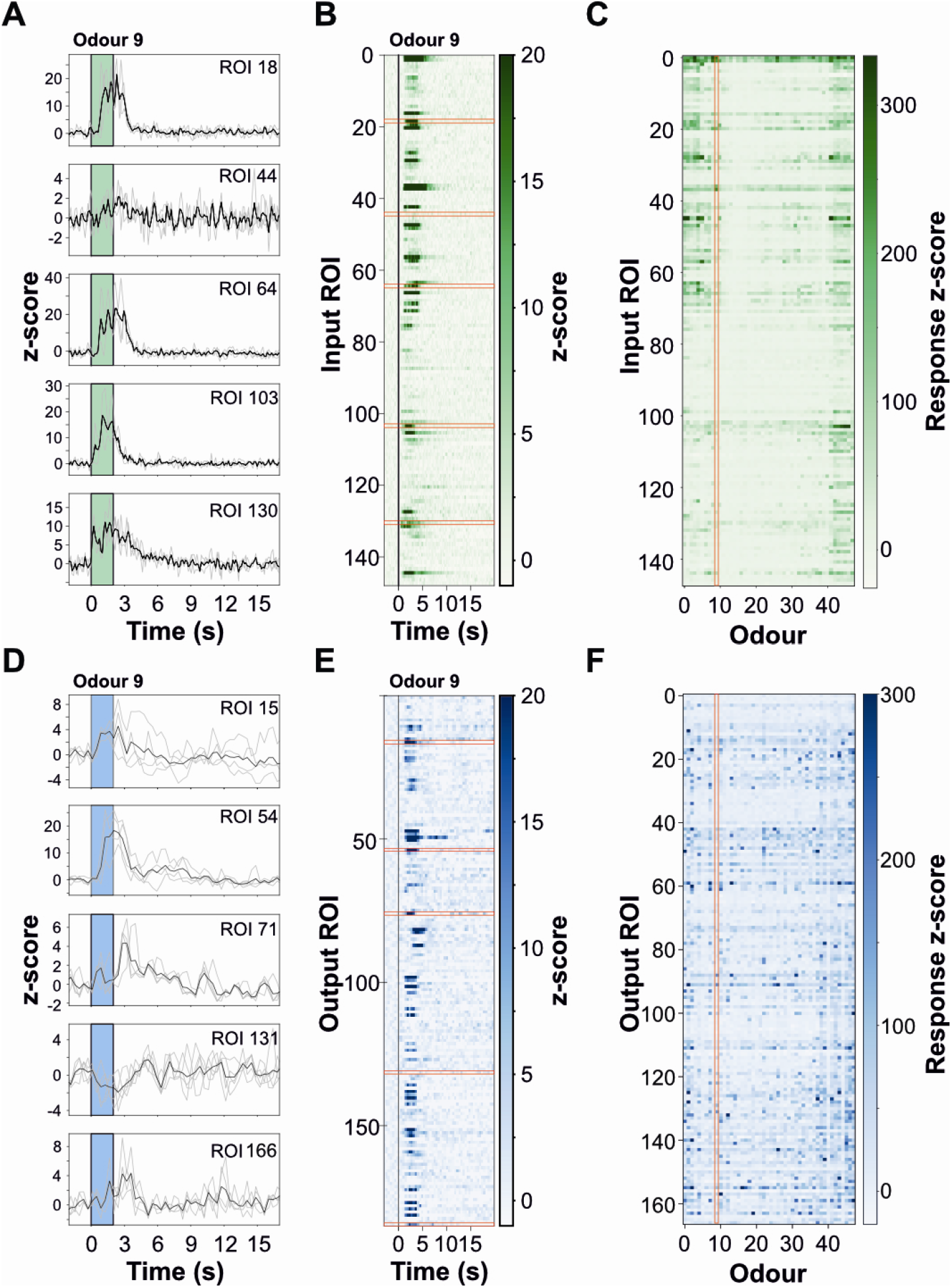
Input and output responses show structured and diverse odour representations. **(A)** Example calcium traces of five glomerular ROIs in response to odour 9 (4-phenyl-2-butanone). Grey: three individual odour presentations, black: mean activation, green: odour presentation window. **(B)** Calcium transients of all ROIs in response to odour 9 shown as colour maps over time. Red boxes mark ROIs shown in (A), n = 148 ROIs from 5 animals. **(C)** Averaged and z-scored integrated responses of all glomeruli to all odours tested (mean of 3 presentations), red box marks odour 9 shown in (A) and (B). **(D-F)** Same as (A-C) but for glomerular output ROIs. Grey: 3-5 individual odour presentations, black: mean activation, blue: odour presentation window, n = 167 ROIs from 12 animals.

The precise glomerular identity of most recorded regions of interest (ROIs) was unknown, limiting the ability to analyse input–output transformations at the level of individual glomeruli. However, the use of triple-transgenic mouse lines enabled us to monitor calcium signals from both input and output in one genetically identified glomerulus, M72. This glomerulus exhibited highly similar activation patterns between input and output across most odours (normalised response difference: 13.46% ± 13.17%; mean ± s.d., *p* = 0.92), but also showed striking differences for a subset of stimuli (**Figure S2.1;** (Arneodo et al., 2018)).

Together, these data provide detailed odour response profiles at both the OB sensory input and output levels. This high-dimensional dataset forms the foundation for analysing how odour representations are structured and transformed within the bulb. To understand how these structured responses differ between input and output stages, we next examined the correlation and the dimensionality of the respective odour representations.

### Odour representations become decorrelated and increase dimensionality from input to output

To understand how the structure of glomerular response patterns differs between input and output stages, we next asked how odour representations differ in their pairwise correlations. The correlation level of response patterns is a critical parameter for the reliable classification of sensory input (Bishop, 1995; Moreno-Bote et al., 2014). Decorrelation or even whitening of sensory input has been described in the visual (Pitkow & Meister, 2012) and auditory system (Saberi & Petrosyan, 2005). In olfaction, decorrelation of odour representations from input to output has been observed in multiple species including locusts (Stopfer et al., 2003), drosophila (Wilson & Laurent, 2005), zebrafish larvae (Friedrich & Laurent, 2001; Wanner & Friedrich, 2020), and mice (Gschwend et al., 2015). Whitening of activity patterns in the zebrafish olfactory bulb is mediated by local inhibitory interneurons and is thought to facilitate pattern classification (Wanner & Friedrich, 2020). Here, rather than simply confirming that decorrelation occurs, we investigated the structure and degree of correlation change from input to output in the mouse OB, and sought to quantify the nature of this transformation.

The response profiles of glomerular ROIs for sensory input (n = 148) and output (n = 167) across the odour panel (n = 47 odours) were used to calculate *pattern correlations* at the input and output, defined as the correlation of population responses to different odour pairs. For the input correlation matrix, a block-like clustering and an overall higher correlation became visible (**Figure 3A, left**) that remained largely absent for the output correlation matrix (**Figure 3A, right**), indicating decorrelation. This decorrelation from input to output was consistent across trials (**Figure S3.1A**) and did not change markedly when ordering odours based on hierarchical clustering of the output data (**Figure S3.1B**,**C**).

**Figure 3:**
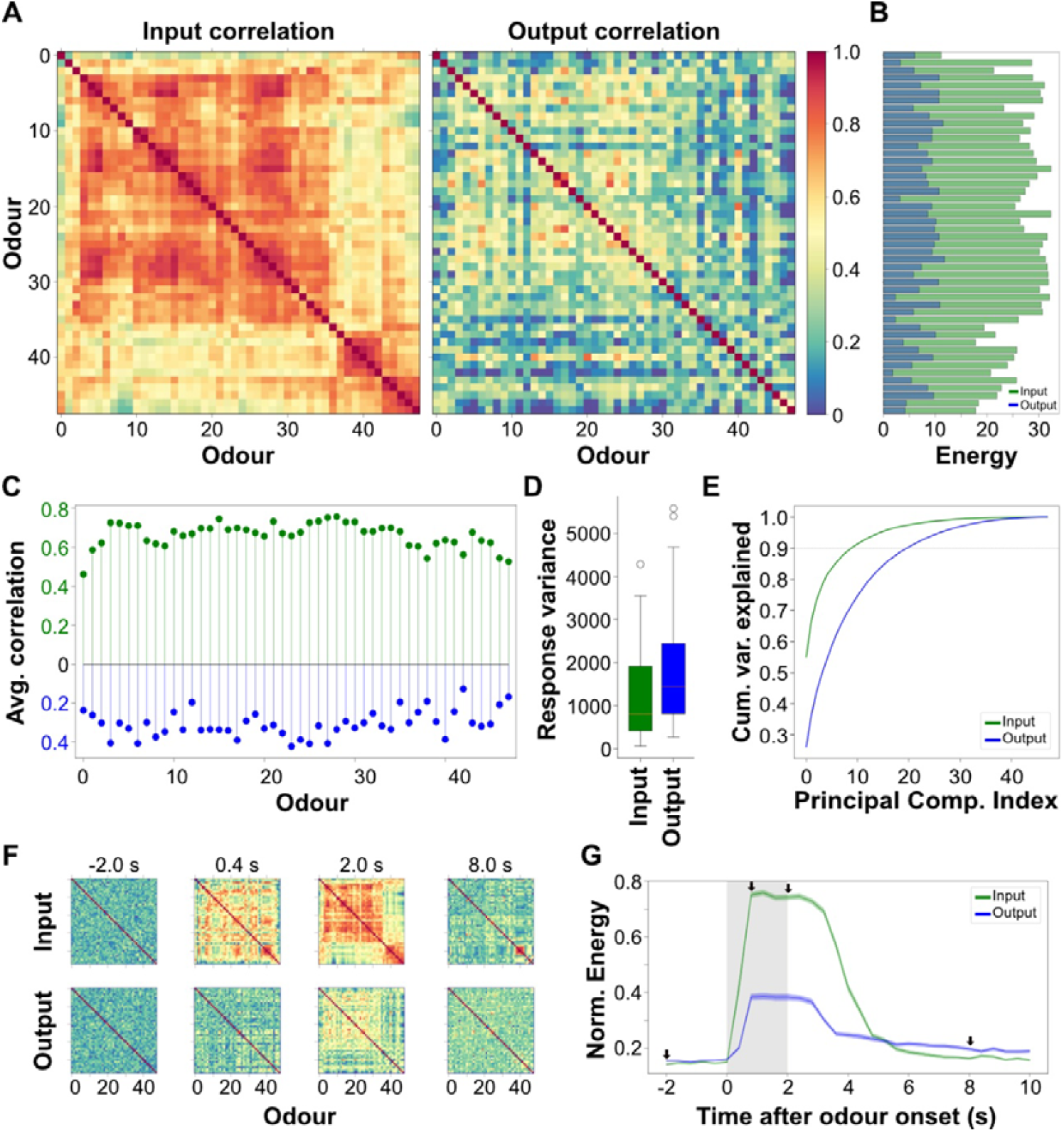
Odour representations become decorrelated from input to output. **(A)** Odour correlation maps of all ROIs for sensory input (left) and output (right). The odour sequence was determined by hierarchical clustering based on the Euclidean distance of the input data and applied to both matrices for comparability (see Methods). **(B)** Odour correlations plotted as Energy (summed correlations squared minus 1, Methods). Input: green, output: blue. **(C)** Mean pairwise correlation across all odours for input (green) and output (blue) signals. For clarity of direct visual comparison, the values for output correlation are plotted below the x axis. **(D)** Response variance across odours, averaged across glomeruli estimated from covariance matrices of sensory input and output (input: 1308 ± 1097; output: 1737 ± 1278; mean ± s.d.; *p* = 0.08), showing no significant difference. **(E)** Cumulative variance explained by the principal components of the input and output responses. Output responses require more components to capture the same proportion of variance (crossing 90% threshold for Input: 10 PCs, Output: 21 PCs) indicating increased dimensionality. **(F)** Example time-resolved correlation matrices for input (top row) and output (bottom row), computed in 400 ms time windows across a trial. Time point 0 marks the onset of odour presentation. **(G)** Correlation energy of input (green) and output (blue) responses plotted over time. Arrows indicate time points corresponding to the matrices shown in (F). Grey shaded area: odour stimulation period. Data were recorded using 47 odours; input: *n* = 148 ROIs from 5 animals; output: *n* = 167 ROIs from 12 animals.

To quantify correlation strength, we calculated the correlation energy for each odour (squared sum of pairwise correlation coefficients, subtracting 1 to remove self-correlation; **Figure 3B**). Correlation energy was significantly lower in the output compared to the input (input: 26.79 ± 4.69; output: 7.70 ± 2.68; mean ± s.d.; p = 3.42e-42), and the majority of odours showed decreased average pairwise correlations at the output level (**Figure 3C**), confirming a broad decorrelation of odour representations. Importantly, this occurred without a significant reduction in overall response variance (**Figure 3D**), indicating that the transformation decorrelates but does not whiten the input signals. To compare the dimensionality of the input and output datasets, we computed the variance explained by the principal components and plotted the cumulative sum of these variances, normalised by the total variance **(Figure 3E)**. The cumulative variance for the input data reached its maximum value more rapidly than that of the output data. This indicates that fewer principal components were needed to explain most of the variance in the input dataset (10 PCs explaining 90% of the variance) compared to the output (21 PCs). In other words, odour representations at the input level occupied a lower-dimensional subspace than those at the output level.

To assess the temporal dynamics of this transformation, we computed correlation maps in 400 ms windows across the trial duration. Output correlations remained consistently lower and less structured than input correlations throughout the trial (**Figure 3F**). Correlation energy rose rapidly at odour onset but remained elevated for the input signal compared to the output (**Figure 3G**). As a further control, we investigated whether the decorrelation observed in the output data was not due to noise level by introducing random noise to the output data. To test this, we computed the correlation between even and odd trials and found a significant auto-correlation, ruling out the possibility that decorrelation was simply due to noise (**Figure S3.1D**). We further confirmed these findings at the level of mitral/tufted cell somata, where odour correlations similarly showed reduced structure and lower overall correlation compared to input, and could not be explained by subsampling or noise (**Figure S3.2**). Comparable correlation patterns were also observed in recordings from awake, head-fixed mice, indicating that the reduced correlation structure of mitral/tufted cell responses is preserved across behavioural states (**Figure S3.2**). To examine whether this transformation can be observed for specific glomeruli, we identified 17 input–output ROI pairs centred around the M72 glomerulus that were matched in both spatial position and odour tuning (**Figure S3.3**). Analysing the response structure of these matched glomeruli revealed that output responses were also less correlated, spanned a higher-dimensional space, and required more principal components to explain the same variance (**Figure S3.4**), supporting the idea that OB processing expands and decorrelates odour representations while preserving functional identity.

Together, these results show that odour representations in the mouse OB become decorrelated from input to output, while preserving overall pattern variance. This transformation could enhance the separability of odour representations and may support improved decoding performance. We next examined whether the observed decorrelation leads to enhanced classification of odour identity from glomerular activity.

### Odour identity is more accurately decoded from output than input activity

Building on our observation that odour representations are decorrelated from input to output, we next asked whether this transformation improves the discriminability of odour identity based on glomerular activity.

To assess how classifier performance scaled with the number of glomerular ROIs, we trained linear classifiers on odour-evoked activity profiles integrated over a 5-second window from odour onset using increasing subsets of randomly selected ROIs (see Methods). As a control, we repeated this analysis with shuffled odour labels. Classifier accuracy increased with the number of ROIs and remained well above chance for both input and output data (**Figure 4A,B**). Output ROIs were substantially more informative than input ROIs: accuracy saturated at around 60 of the 167 available output ROIs, while no such saturation was observed for the full set of input ROIs. When we trained the same linear model on all glomeruli from the input data, odour identity could be correctly predicted from among 48 stimuli (47 odours and mineral oil as blank) with an accuracy of 93.4 ± 1.0% (mean ± s.d.; chance level: 1/48 ≈ 2.083%). Even higher accuracy was achieved with the output data, where mean performance reached 99.5 ± 0.4%).

**Figure 4.**
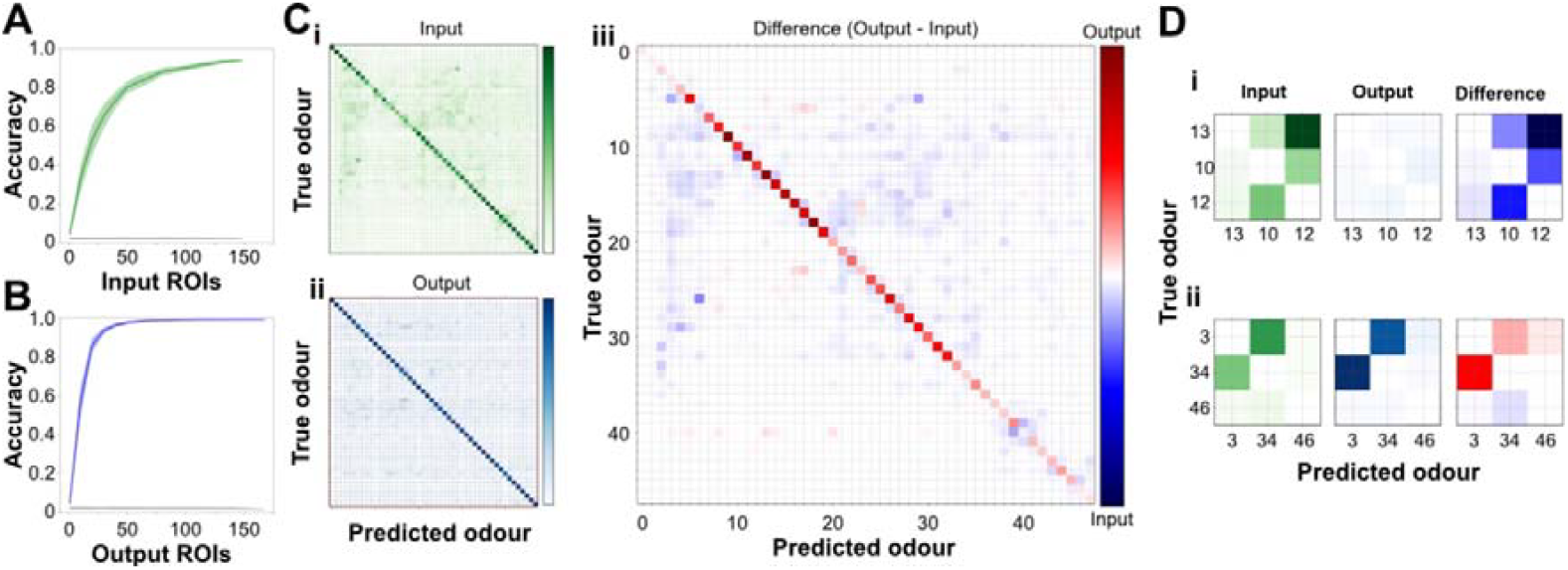
Odour identity is more accurately decoded from output than input activity. **(A)** Classification accuracy of a set of LDA classifiers trained on input calcium responses to 47 odours, plotted as a function of the number of ROIs included. Shown are mean ± s.d. across 50 repetitions (up to 148 glomeruli from 5 individual animals; green: input, black: shuffle control). **(B)** Same as (A), but for the output data (mean ± s.d. across 50 repetitions; up to 167 glomeruli from 12 individual animals; blue: output, black: shuffle control). **(C)** Confusion matrices showing LDA-based classification results averaged across 100 iterations with 30 randomly selected glomeruli per iteration, for (i) the sensory input, (ii) the output, and (iii) the difference between input and output (output - input). Labels predicted by the classifier are shown on the x-axis and true labels on the y-axis. Red indicates higher output misclassification; blue indicates higher input misclassification. **(D)** Confusion submatrices showing selected odours with exclusive mismatch patterns. (i) Three odours with selective input mismatches (high input confusion, accurate output classification). (ii) Three odours with selective output mismatches (accurate input classification, high output confusion). Diagonal elements (correct classifications) are masked to highlight off-diagonal mismatches.

To evaluate how well individual odours could be predicted based on OB input and output signals, we performed a classification analysis across all tested odours. To ensure robust estimates, we repeated the classification 100 times each using responses from 30 randomly selected glomeruli (mean accuracies ± s.d. across iterations for input: 65.1 ± 6.0%, output: 93.3 ± 1.8%). We computed averaged confusion matrices for input and output signals from these iterations, capturing prediction accuracy across the full odour set (**Figure 4C**). While certain odours are predicted more accurately from the output and others from the input, the overall classification accuracy is consistently higher for the output signals. The difference matrix (output minus input) reflects correctly classified odours, while off-diagonal entries indicate misclassifications. Red colours off-diagonal denote higher misclassification rates for output and blue for input. Notably, some odours were predicted more accurately from the output than the input, suggesting transformation of odour representation and a possible enhancement in the OB, while others showed the opposite pattern. To systematically identify clear examples of these opposing transformations, we applied exclusive mismatch criteria: selecting odours with high misclassification on one side (input or output) but accurate classification on the other (**Figure 4D**). Three automatically selected odours exhibit selective input mismatches - poor input classification (off-diagonal green) but accurate output predictions (diagonal blue) (**Figure 4Di**). Conversely, other odours show the opposite pattern: accurate input classification but degraded output discrimination (**Figure 4Dii**). These contrasting examples demonstrate that OB transformations can either sharpen or degrade odour separability, with the overall population effect favouring enhanced discrimination.

Taken together, these results demonstrate that odour identity can be reliably decoded from glomerular responses, and that the output representations support significantly better classification performance than the input. This enhanced decoding is consistent with the observed decorrelation and suggests that the input–output transformation in the OB increases odour separability in a way that benefits downstream readout. Further, these results indicate that OB processing selectively enhances the discriminability of some odours while reducing that of others, reflecting a transformation rather than a uniform gain in odour information.

While these analyses reveal that decorrelation improves decoding, they do not explain how the olfactory bulb achieves such a transformation mechanistically. To address this, we next turned to computational modelling, using the recorded inputs to constrain candidate connectivity motifs.

### Relating decoding accuracy to decorrelation

Earlier we showed that odour responses are decorrelated at the output of the OB relative to the input. We have just shown that decoding accuracy is higher at the system output than at the input. Are these two findings related? This question is complicated because accuracy is determined not only by the correlations of the mean responses (signal correlations), but by the scatter of the single-trial responses relative to the odour-specific means (noise correlations).

We answered this question by taking a geometric approach. From this perspective, the correlations of the trial-averaged odour response vectors reflect the angles between these vectors. In fact, when response vectors all have zero mean (across ROIs), the Pearson correlation of a pair of response vectors is the cosine of the angle between them. Our data do not have zero mean across ROIs, so the Pearson correlations and cosine correlations are in general different. However, the two sets of values remain highly correlated (see **Figure S5.1**), and we used cosine correlations below as it simplified the analysis.

Using the cosine of the angle between response vectors as our measure of correlation, we reasoned that the contribution of correlations to the increased accuracy achieved from input to output would be the improvement in accuracy achieved when the input data was modified to have the same correlations as the output, while keeping all other aspects of its encoding geometry unchanged. The projection of this procedure into the span of two odour responses is shown in the first two panels of **Figure 5Ai**. The accuracy before this transformation was ~70%, and increased to ~100% afterwards (**Figure 5B**, solid orange arrow), even higher than its value of ~90% at the output (**Figure 5B**, grey point). Reasoning similarly but from the output side, we performed the same procedure on the output responses, transforming them to have the same correlations as the input responses while keeping their remaining geometry unchanged. The projection of this procedure in the span of the same two odour responses as in **Figure 5Ai** is shown in the final two panels of **Figure 5Aii**. When we transformed the output in this way we observed a decrease of decoding accuracy from ~90% to ~60% (**Figure 5B**, dashed orange arrow), lower than the value of ~70% observed at the input (**Figure 5B**, grey point). By performing further transformations, we were able to sequentially transform the input dataset to the output dataset and vice versa, computing the decoding accuracy after each step. The remaining panels of **Figure 5A** show these steps projected into the span of two odour means, and **Figure 5B** shows the corresponding accuracies. Interestingly, the resulting changes in accuracy were not monotonic (**Figure 5B**). Decoding accuracy also depended on the order in which the transformations were performed (**Figure S5.2, S5.3)**, so that the contribution of matching correlations could be both greater than, and less than, what we observed when performing this step first. Nevertheless, when the only modification to a dataset was to match its correlations to the other, we observed an absolute change in accuracy of ~30%, which we took to be the contribution of correlations to the decoding accuracy.

**Figure 5:**
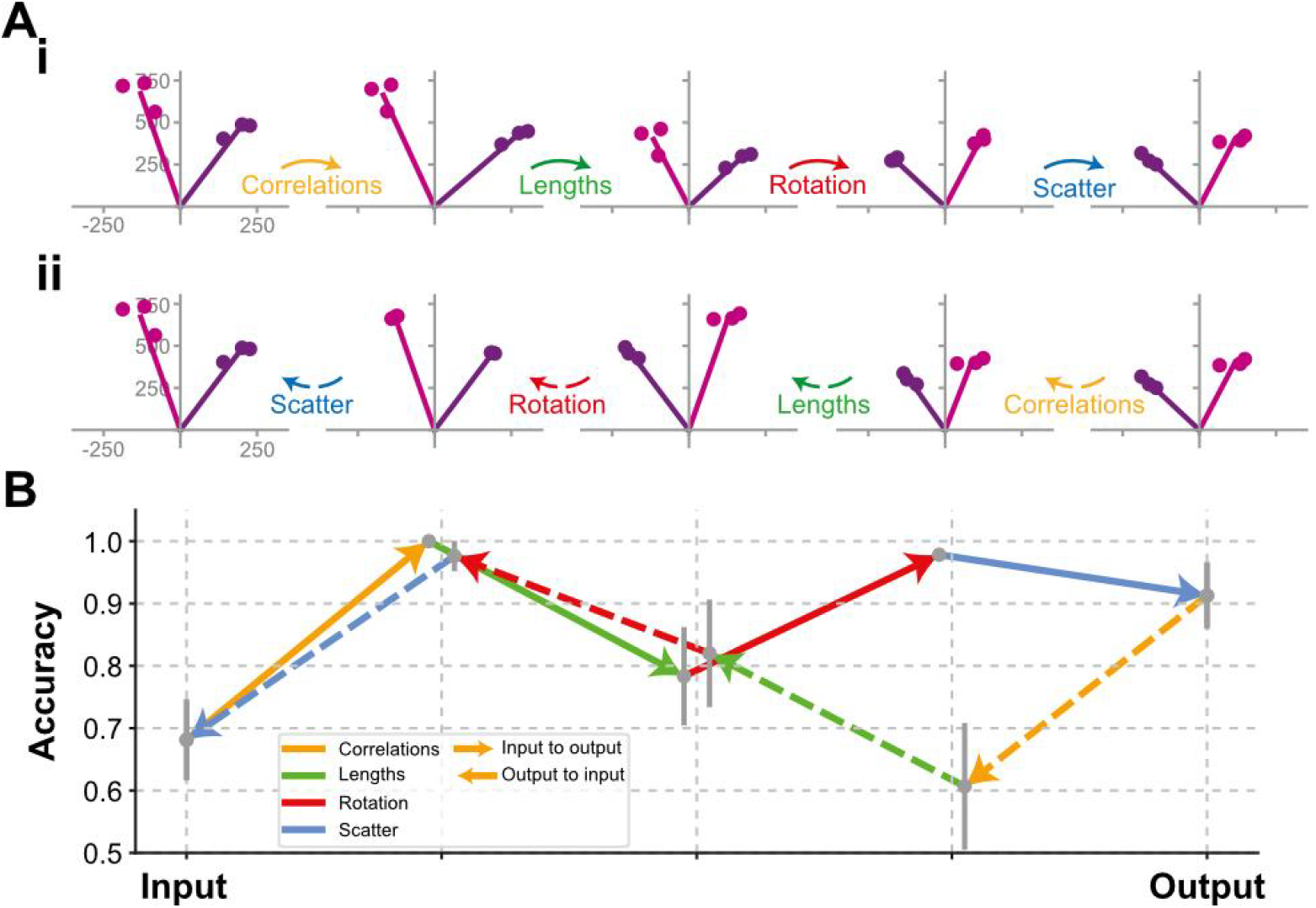
Relating accuracy to decorrelation. The contribution of decorrelation to the decoding accuracy is determined by transforming the input responses into output responses, and vice versa, through a sequence of linear transformations. **(Ai)** Projection of the transformation converting input to output into the span of the mean responses to two odours (pink and purple lines). Points are the projections of the three trials available for each odour. Transformations are indicated below the arrows linking plots. The first transformation matches the cosine correlations between the mean responses to those of the output. The second matches the lengths. The third rotates/reflects responses to align with the output. The final transformation produces an exact match to the output by adjusting the position of the individual trials relative to the odour-specific means. See Methods for details. **(Aii)** As in (Ai) but transforming the output into the input. **(B)** Mean (grey points) ± 3 s.d. (grey lines) of the accuracy at each stage of the transformation described in (Ai,ii). Coloured lines linking points indicate the transformation performed. Arrows and line styles indicate the directions of transformation. Points between input and output have been staggered for clarity.

### Understanding the input–output transformation using constrained connectivity

To understand how the OB might implement the experimentally observed decorrelation, we constructed a simple linear model consisting of glomerular inputs, mitral cells, and inhibitory connectivity between them (**Figure 6A, 7A**). This allowed us to test which classes of connectivity could reproduce the transformation we measured. Just as in the real circuit, each model mitral cell was excited by a single parent glomerulus. It was also inhibited to varying degrees by all other mitral cells. The strength of this inhibition was determined by a connectivity matrix whose elements represented the effective inhibition of each mitral cell by every other and implemented disynaptically by granule cells in the real circuit (and to a possibly lesser extent by glomerular and external plexiform layer interneurons).

**Figure 6:**
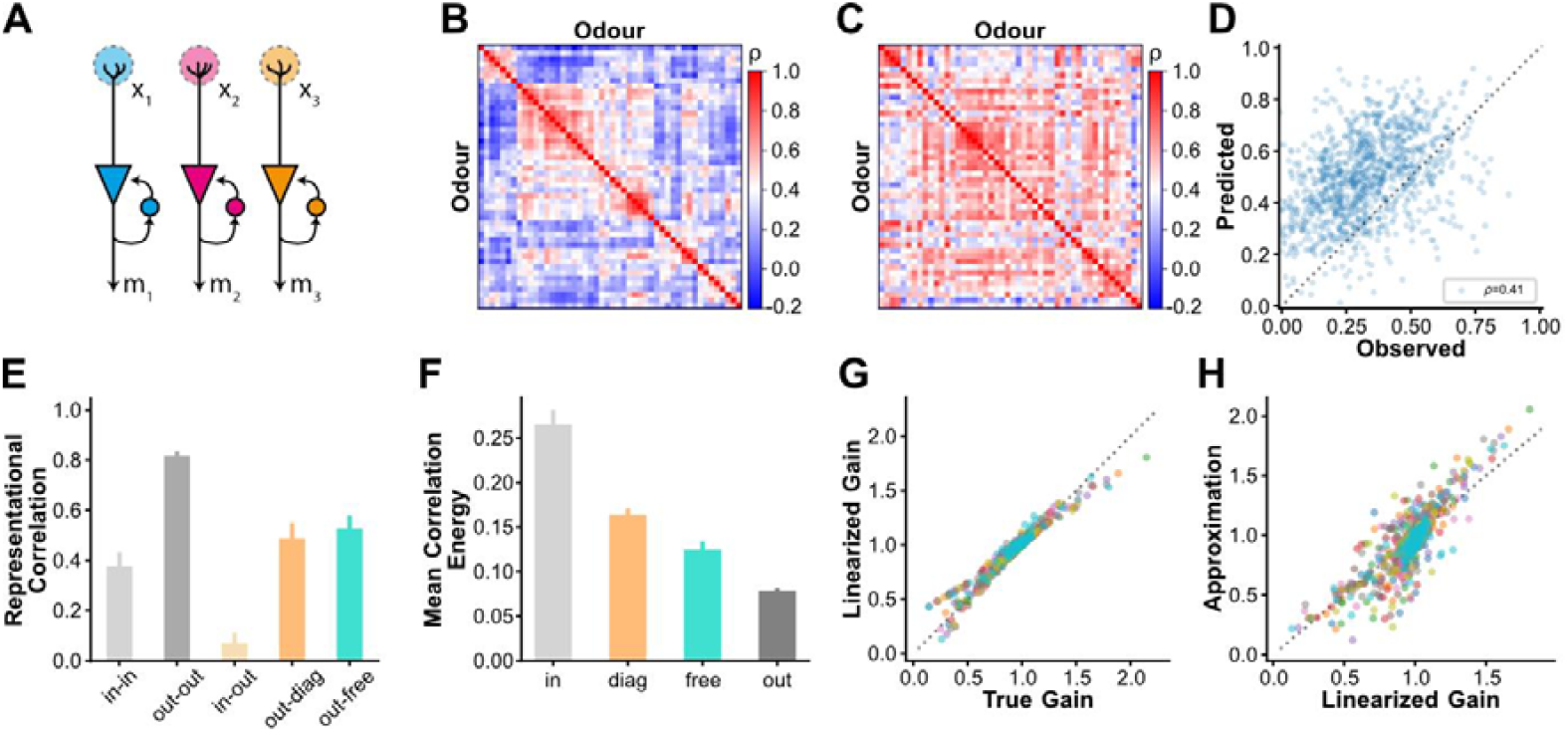
Fitting without lateral connectivity. **(A)** Circuit schematic of the “Diagonal” model, showing the transformation of three input channels (colours). No lateral interactions are allowed, leaving only self-excitation or inhibition as the possible transformations. **(B)** Pearson correlations of trial-averaged glomerular responses. **(C)** Predicted Pearson correlations when trained to produce the data in (B) from the corresponding input data, using the regularization value determined by cross-validation on single trials. **(D)** Observed vs. predicted correlations for every pair of odours (each element above the diagonals of B vs. C). The legend indicates the overall Pearson correlation of the observed and predicted data. **(E)** Representational correlations. A representation is defined as the Pearson correlations of responses to all unique pairs of odours i.e. the above-diagonal elements of panels B and C. The panel shows the mean (bars) + s.d. (whiskers) computed over 10 trials of the Pearson correlations of the representations of different datasets. *“in-in”*: one random sample of the input responses against another; *“out-out”*: one random sample of the output responses against another; *“in-out”*: one random sample of the input responses against a random sample of the output responses; *“out-diag”*: one random sample of the output responses against the predictions of the diagonal model that was trained on a different subset of trials. *“out-free”*: as in *“out-diag”* but for the “Free” model, described below. **(F)** Mean (bars) + s.d. (whiskers) of the mean energy of the off-diagonal elements of the representations each dataset. *“in”*: input responses, 30 random samplings of single trials; *“out”*: 30 random samplings of single trials; “diag” & “free”: predictions on 20 random samplings of held out trials for each model. **(G)** The true values of the gains applied to each unit (dots) on 10 different trials (colours) vs. their linearizations around 1. **(H)** The linearized values of the gains in panel (G) vs. their approximations.

When an odour is presented to such a circuit, the glomerular activations drive the mitral cells to an odour-specific steady-state, which we took to be the output of the model. The interactions among mitral cells mediated by the connectivity mean that this output will in general be different from the input. However, the simplicity of our model means the relationship is linear, with an implicit feedforward connectivity determined by the recurrent weights by which mitral cells inhibit each other. In what follows, we will work primarily with the connectivity expressed in this feedforward form.

We next searched for feedforward connectivity that made the model outputs as close as possible to those experimentally recorded. One way to do this would be to compare modelled and experimental outputs directly. That is, if we knew the identities of the input and output glomeruli, we could predict how connectivity transforms the inputs to an identified set of glomeruli to corresponding outputs. We could then compare those predictions to the experimental recordings from the same glomeruli, and adjust the connectivity to reduce the discrepancy.

However, we only knew the identity of a few glomeruli with certainty. Therefore, rather than comparing the responses of individual glomeruli to each odour, we compared the similarity of the population responses for all pairs of odours. We computed the similarity of a pair of stimuli as the Pearson correlation of the corresponding population response vectors. Our model’s predicted similarity was computed in the same way, by first using the connectivity to transform each recorded input to a predicted output, and then computing the correlation between two such outputs.

Fitting Pearson correlations is numerically straightforward but interpreting the resulting connectivities analytically can be complex. Therefore, during our fitting procedure we matched covariances instead. Since Pearson correlations are derived from covariances, perfectly matching covariances would automatically match Pearson correlations as well. We computed the difference between the observed and predicted representations by summing the squared difference between the covariances over all stimulus pairs and adjusted the connectivity to reduce this difference. Because we frequently had more connectivity parameters (i.e. synapses) than representation parameters to fit, we regularised the connectivity to prefer small values in its equivalent lateral connectivity form. Such regularisation preferred to pass through the input to the output unchanged. We controlled the strength of this regularization through a parameter whose value we optimized using cross-validation. In practice, this meant that the fitted connectivity reflected only the minimal adjustments to the input necessary to reproduce the observed output.

### Decorrelation without lateral connections

To better understand the transformations different connectivities can implement, we tested connectivity from two families spanning the range of possibilities. We began by fitting connectivities that lacked lateral connections entirely. Because the resulting connectivity matrices would only have terms on the diagonal, indicating self-connections, we called these “Diagonal” models. We describe these below. After testing the Diagonal models, we removed all connectivity constraints and fit the resulting “Free” model. We describe those results in the next section.

In the model without lateral interactions, the only possible transformation was self-amplification or self-inhibition (see schematic in **Figure 6A)**. Given the anatomical prominence of lateral connectivity in the OB, we expected that it would be necessary for decorrelation. Surprisingly, however, we found that this simple model was able to perform nearly as well as the “Free” model which also allowed lateral connections (see below). We trained the model by first selecting a random subset of trials and then adjusting the gains applied to the input responses on those trials so that the corresponding representations best matched those of the recorded output responses for those trials. We tested the model’s performance by using the resulting channel gains to perform the input–output transformation on a different, unseen, set of trials. We repeated this procedure 10 times and used cross-validation to set the value of a regularization parameter (see Methods).

The observed output correlation structure for trial-averaged output data is shown in **Figure 6B** alongside the model prediction (**Figure 6C**), plotted pairwise in **Figure 6D**. Although the predicted correlations were typically higher than those observed, as apparent in the reddish hue of **Figure 6C** vs. **Figure 6B** and the vertical shift of the points in **Figure 6D**, predicted and observed pairwise odour similarities were strongly aligned, as illustrated by their correspondence across all odour pairs. Across repeated random samplings, the model produced representations that were as strongly correlated with the output as those of the Free model, and substantially more so than the original input representations (**Figure 6E**, ‘in-out’).

Because linear correlation is insensitive to uniform shifts in similarity values, we additionally quantified correlation structure using the mean correlation energy, defined as the average squared correlation across all odour pairs. This metric revealed that the diagonal model captured a substantial fraction of the decorrelation between input and output representations, despite lacking lateral interactions (**Figure 6F)**. Specifically, the model accounted for approximately half of the total reduction in correlation energy observed experimentally, indicating that channel-wise gain modulation alone contributes significantly to decorrelation.

Staying momentarily with this simple model, we next asked what determines whether a channel is amplified or suppressed. Fitting covariances can introduce nonlinearities that can make interpreting the learned connectivities difficult. However, we found that the true gains learned by the model were well captured by their linearizations around their regularization value of 1 (**Figure 6G**). As we detail in the Methods, the linearized gains are determined by two factors. First is how much each unit’s own representation, defined as the product of its responses to each odour pair, overlaps with the difference in representations between the output and input. The second is how much the unit’s representation overlaps with that of the other units, weighted by the gains applied to those units. The need to account for the gain applied to the other units in this determination complicates interpretation. Remarkably, we found that in determining the gains applied to each unit, we could approximate the influence of the other units as a scalar multiple of their mean representations **(Figure 6H)**. This meant that we could approximate the gain applied to each using only knowledge of the input and output representations, rather than also the gains applied to every other unit.

In summary, partial decorrelation can be performed without lateral connectivity, by adjusting the gains of the input channels. These gains are determined by the overlap of each unit with the representational difference of the input and output, plus a constant multiple units average overlap with all the other units.

### Free connectivity

Having shown that channel-specific scaling alone can account for a significant fraction of the decorrelation from input to output, we next asked whether incorporating lateral connectivity could further improve the match to the experimental data. Anatomically, the OB contains extensive lateral interactions mediated by inhibitory interneurons, raising the question of how much these contribute to the observed decorrelation. To address this, we relaxed the constraints on the model and allowed connectivity to be learned freely (**Figure 7A**).

**Figure 7:**
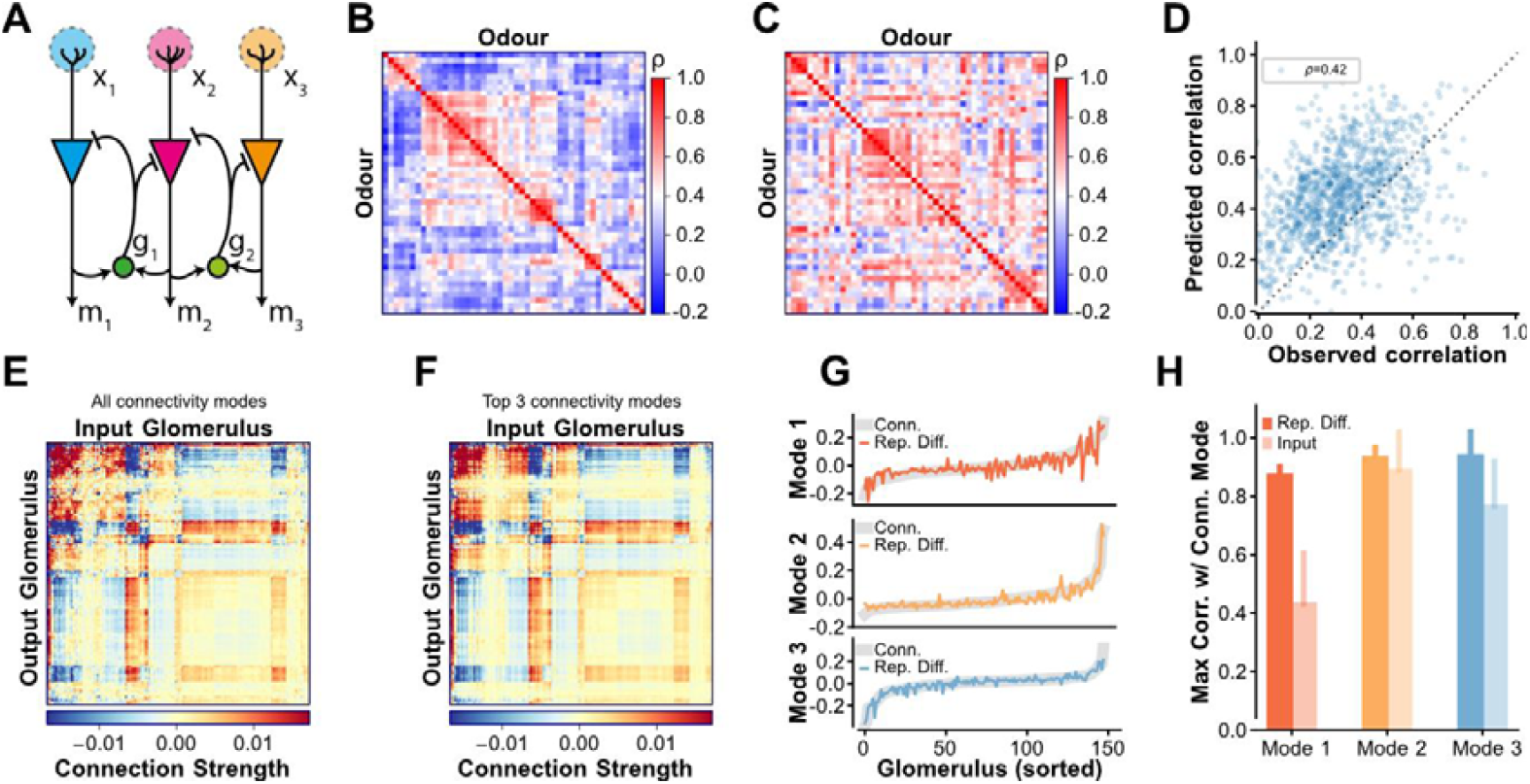
Fitting lateral connectivity to the input–output transformation. **(A)** Circuit schematic, showing three glomerular input channels, *x*_1_, *x*_2_, and *x*_3_ (colours) are transformed into mitral cell outputs *m1, m2*, and m3, by lateral interactions mediated by granule cells *g*_1_ and *g*_2_. Instead of fitting the lateral connectivity, we fit the corresponding effective feedforward connectivity from input to output channels. **(B)** Pearson correlations of trial-averaged glomerular responses. **(C)** Predicted Pearson correlations when trained to produce the data in (B) from the corresponding input data, using the regularization value determined by cross-validation on single trials. **(D)** Observed vs. predicted correlations for every pair of odours (each element above the diagonals of B vs. C). The legend indicates the overall Pearson correlation of the observed and predicted data. **(E)** Effective feedforward connectivity used to make the prediction in panel C. **(F)** Approximation of the connectivity in panel (E) using only the top 3 modes. **(G)** Agreement of the top 3 connectivity modes (grey) with the best-matching mode of the representational difference (colours). **(H)** Mean (bars) and s.d. (whiskers) of the highest correlation, measured as scalar product, of each connectivity mode with the representational difference modes (dark) or input modes (light), computed over 10 trials.

With appropriate regularization determined by cross-validation, this model could reduce input–output correlations by approximately 75% (**Figure 6F**). The output of this model when trained on the trial-averaged inputs to match the trial-averaged output representation (**Figure 7B**, same as in 6B) using the regularization value found by cross-validation, is shown in **Figure 7C**. The close element-by-element correspondence between predicted and observed values highlights this match (**Figure 7D**).

We next investigated the logic of the connectivity that produced this match. Because we did not impose additional constraints on the connectivity beyond regularization, the learned connectivity could be complex, making understanding difficult. Fortunately, we found the learned connectivity to be highly structured. An example of the effective glomerulus-to-glomerulus feedforward connectivity is shown in **Figure 7E**. Most obviously, this connectivity was symmetric. It was also well approximated by its three top connectivity modes (**Figure 7F)**.

As we show in the Methods, our connectivity solutions are symmetric because we select them by how well they produce the output representation, while subjecting them to regularization. Importantly, representations reflect the similarity of pairs of responses rather than the responses themselves. These are in turn determined not by how inputs are sampled by each output unit, as tabulated in the feedforward connectivity, which may or may not be symmetric, but rather by the similarities in how inputs feed forward to outputs. The symmetry of these similarities, combined with the regularization pressure on the feedforward connectivity that pulls it towards the identity, which is also symmetric, ultimately produces a symmetric connectivity solution.

Thus, while the true connectivity may or may not be symmetric, the solutions we recover reflect the structure of the representational transformation rather than the precise wiring of the circuit. This highlights that only balanced, low-dimensional interactions are required to explain the data, even if biological circuits implement them through asymmetric connections.

As mentioned in the previous section, matching covariances introduces nonlinearities that can make interpreting the resulting connectivity difficult. When performing our fits, we found that we needed to use a large amount of regularization. Such high regularization meant that the learned values did not deviate significantly from their regularization targets, allowing us to approximate the connectivity solutions by linearization. We found that the top three connectivity modes were well captured by the top three modes of the representational difference between input and output, though not necessarily in the same order (**Figure 7G**). Importantly the agreement with these modes was better than the agreement with those of the input alone, as would be predicted if the outputs were fully decorrelated (**Figure 7H**).

## Discussion

Decorrelating transformations of sensory input have been reported in the olfactory bulb (OB) of mice and other species including zebrafish and flies, and are often attributed to inhibitory interactions that whiten or orthogonalize population responses (Bhandawat et al., 2007; Friedrich & Laurent, 2001, 2004; Gschwend et al., 2015; Lin et al., 2014; Otazu et al., 2015; Stopfer et al., 2003; Wanner & Friedrich, 2020; Wilson & Laurent, 2005). However, most prior studies were based on relatively small stimulus sets or focused exclusively on output activity, leaving open how systematic and general such transformations are in the mouse OB, and what minimal circuit motifs are sufficient to generate them. Our goal was to address these gaps by comparing glomerular input and output responses to a large and chemically diverse odour panel, and by coupling these measurements with mechanistic modelling. This approach allowed us not only to confirm that odour representations decorrelate from input to output, but to quantify the transformation across a broad stimulus space and to identify the connectivity rules that can account for it.

To characterise input and output response profiles, we recorded summed glomerular signals using two-photon calcium imaging in OMP-GCaMP6f and Tbet-GCaMP6f transgenic lines. This allowed us to capture OSN input signals and bulk M/T cell output signals in matched dorsal OB fields of view, albeit not in the same animals (Ackels et al., 2020, 2021; Bosch et al., 2022). The stable representation of landmark glomeruli across animals, thus, provided a basis for comparative analysis. Consistent across animals, glomerular responses showed reliable activation patterns, validating this approach as a population-level assay of input– output transformations. Importantly, we also imaged activity in identified mitral and tufted cell somata, where pairwise correlations were similarly reduced compared to the summed dendritic signals within glomeruli. Recordings from awake, head-fixed mice revealed a comparable correlation structure, indicating that the observed decorrelation is not dependent on the animal’s state (**Figure S3.2**). While input and output were not recorded simultaneously in the same preparation, the reproducibility of glomerular maps across animals supports the validity of this comparative strategy. Future dual-colour imaging approaches (Storace & Cohen, 2017) may further refine these analyses by enabling direct within-animal comparisons.

Our experiments revealed a robust decorrelation of odour representations from input to output across 47 diverse odours. Importantly, this decorrelation was not accompanied by a loss of overall variance, indicating that the transformation reduces correlations without whitening the input (Moreno-Bote et al., 2014; Olshausen & Field, 1996; Pitkow & Meister, 2012; Simoncelli & Olshausen, 2001; Wanner & Friedrich, 2020). As shown in **Figure 3**, output representations occupied a higher-dimensional subspace than inputs, consistent with an expansion of coding capacity. These results extend prior reports of decorrelation in the mouse OB (Gschwend et al., 2015; Otazu et al., 2015) by demonstrating that the effect is systematic across a large stimulus set and can be quantified in terms of dimensionality expansion as well as correlation structure.

Correlations in the OB changed dynamically over the course of a trial: both input and output activity became more correlated shortly after stimulus onset and then progressively decorrelated, with outputs maintaining lower correlation levels than inputs throughout (**Figure 3**). This temporal progression is consistent with previous work showing that decorrelation continues to develop after stimulus onset and enhances discrimination (Friedrich & Laurent, 2001, 2004; Wanner & Friedrich, 2020). Local inhibitory networks can contribute to these dynamics by shaping correlations on distinct timescales (Giridhar et al., 2011; Kato et al., 2012), and cortical feedback is also known to modulate bulb activity (Boyd et al., 2015; Chae et al., 2022). Such decorrelation is typically performed using lateral connectivity. Indeed, our linear model that used lateral connectivity was best at reproducing the input–output transformation. Interestingly, our modelling showed that the bulk of the transformation achieved by the lateral connectivity could be achieved using simple feedforward gain modulation. This suggests that inhibitory and feedback influences may refine the temporal structure of population activity, but a significant driver of robust decorrelation across our large odour panel is feedforward scaling of glomerular channels. In natural environments, odours are encountered not as steady pulses but as spatiotemporally complex plumes (Ackels et al., 2021; Gumaste et al., 2024; Marin et al., 2021; Murlis et al., 2000; Mylne & Mason, 1991), and it will be important in future work to determine how such plume dynamics interact with the feedforward scaling we identify here.

Another extension of our work comes from linking decorrelation directly to decoding. Linear classifiers trained on glomerular responses showed that odour identity could be reliably decoded from both input and output signals, but with markedly higher accuracy at the output (**Figure 4**). Moreover, by manipulating the correlation structure of the data, we quantified how much of this improvement could be explained by decorrelation itself, establishing a direct link between representational geometry and decoding accuracy (**Figure 5**; Miura et al., 2012; Pashkovski et al., 2020). Thus, decorrelation is not only a descriptive feature of OB processing, but directly improves odour discriminability.

We then asked what circuit motifs could account for this transformation. Surprisingly, modelling revealed that diagonal-only connectivity, equivalent to channel-specific gain modulation without lateral terms, reproduced the majority of the experimentally observed decorrelation that our linear model could capture (**Figure 6**). Allowing free lateral connectivity provided only modest additional improvements, and these were captured by a small number of low-rank modes (**Figure 7**). Given the anatomical prominence of interneuron-mediated lateral interactions in the OB (Arevian et al., 2008; Burton, 2017; Fukunaga et al., 2014; Parrish-Aungst et al., 2007; Urban, 2002), this was unexpected.

Theoretical works have also investigated the benefits and mechanisms of decorrelation. These distinguish two types of correlation (Wick et al., 2010). *Pattern* correlations are the correlations of a population’s responses to different stimuli. These measure how the responses to pairs of *stimuli* covary across a neural population. They are what we in this work and others in the works cited above refer to simply as correlations. *Channel* correlations measure how the responses of pairs of *neurons* covary across stimuli. Theoretical investigations have motivated channel decorrelation on the grounds of efficient coding as a means of maximizing information transmission while minimizing energy expenditure (Atick & Redlich, 1993). Other works have shown how pattern decorrelation can be achieved through channel decorrelation (Wick et al., 2010). Further works have addressed the more general problem of matching one set of pattern correlations to another (Pehlevan et al., 2015). All of these works have used adaptive algorithms that rely on lateral connectivity. For example, the first two show how decorrelation can be achieved by using lateral inhibition to invert the channel correlations present in the input. Interestingly, (Wiechert et al., 2010) showed how decorrelation can be achieved using random recurrent connectivity, without requiring adaptation to input statistics. Indeed, we also observed that lateral connectivity was required to achieve the best match between our input and output responses. What was surprising was that much of this match could be achieved without such connectivity, relying only on gain control of the input channels.

Our findings thus suggest that much of the transformation from input to output can be achieved by feedforward scaling of individual glomerular channels, potentially implemented through inhibitory circuit motifs such as feedforward inhibition. This perspective challenges the prevailing view that decorrelation in the mammalian OB depends primarily on more complex inhibitory networks, such as recurrent or lateral inhibition. Instead, it highlights a simpler and scalable mechanism that selectively amplifies informative channels while attenuating redundant ones. Such a rule is computationally efficient, consistent with redundancy reduction strategies observed in other sensory modalities (Bishop, 1995; Carandini & Heeger, 2012; Litwin-Kumar et al., 2017; Moreno-Bote et al., 2014), and could be implemented by local gain control at the level of glomeruli or M/T cells. Nevertheless, channel gain-control is unlikely to be the only source of decorrelation in complex odour environments, since the number of odour pairs to decorrelate increases quadratically with the number of odours, and would quickly exceed the degrees of freedom provided by the channels. For example, the ~1500 olfactory channels in mice would suffice to decorrelate on the order of √1500 ≅ 40 odours, which is likely far fewer than the number that a given mouse encounters in its environment. Therefore, gain-control must operate in tandem with lateral connectivity modes to perform decorrelation. The low-rank lateral modes we identify may thus correspond to broad inhibitory influences that suppress overrepresented patterns and enhance underrepresented ones, complementing the dominant contribution of feedforward scaling.

While our findings provide a new perspective on OB processing, several limitations should be considered. Our main experiments were performed under anaesthesia and input and output recordings were not obtained from the same animal, potentially reducing contributions from cortical feedback and state-dependent modulation, which are known factors shaping OB activity. However, to address this limitation, we also performed a subset of recordings in awake animals, and found that the key transformations identified under anaesthesia, including robust decorrelation of odour representations, were similarly observed in the awake state, supporting the generalizability of our main findings.

A second important limitation is that much of our analysis was performed at the level of bulk glomerular activity patterns, based on population-averaged signals across defined input and output channels. This approach is well suited to reveal transformations in the mean representation carried by glomerular channels, but it may obscure heterogeneity among individual mitral and tufted cells associated with the same glomerulus (Arneodo et al., 2018; Dhawale et al., 2010; Karadas et al., 2026; Schwarz et al., 2018; Zhang et al., 2025). In particular, granule-cell-mediated lateral inhibition and related inhibitory motifs may contribute to diversifying sister-cell responses, a transformation that would be less visible in bulk glomerular measurements but could be important for expanding the dimensionality of OB output at the single-cell level. Thus, our results argue that feedforward scaling can account for a substantial component of the input–output transformation, but they do not exclude additional circuit mechanisms that operate at finer cellular resolution.

Building on this, our results also suggest a broader functional role for inhibitory circuitry in the OB. If a substantial component of decorrelation can be explained by feedforward scaling at the level of glomerular channels, then granule cells and related inhibitory networks may instead contribute primarily to shaping the temporal, contextual, and cell-specific structure of odour representations. For example, granule-cell-mediated dendrodendritic inhibition has been implicated in coordinating activity across mitral and tufted cells, regulating synchrony and spike timing, and dynamically sculpting population responses (Fukunaga et al., 2014). In addition, inhibitory and feedback circuits may support functions such as gain control, pattern completion under noisy conditions, and the integration of top-down signals related to expectation, learning, or behavioural context (Markopoulos et al., 2012; Otazu et al., 2015). From this perspective, feedforward scaling may implement a core transformation that improves separability of odour representations, while inhibitory circuits provide additional flexibility, enabling the OB to dynamically adapt representations to behavioural demands and internal state.

Our analysis was restricted to spatial patterns of trial-averaged activity. Temporal dynamics, which have been implicated in odour decorrelation in other species, were not captured here and may further refine representations on faster timescales (Ackels et al., 2021; Cury & Uchida, 2010; Friedrich & Laurent, 2001, 2004; Shusterman et al., 2011). Finally, our modelling framework was deliberately linear and abstracted away many biological details, including nonlinear dendritic integration, heterogeneity of inhibitory cell types, and plasticity mechanisms. The fact that such a simple model can capture a significant portion of the observed transformation demonstrates the power of feedforward scaling, but it does not preclude additional roles for more complex circuitry in natural contexts.

Viewed from this perspective, the framework also points to clear experimental predictions. If lateral inhibition were selectively diminished, for example by pharmacological blockade of granule cell circuits, we would expect only modest changes in bulk decorrelation if diagonal gain adjustments are indeed the dominant mechanism. This expectation is consistent with prior work showing that granule cell-mediated transmission had only limited effects on odour discrimination behaviour (Abraham et al., 2010; Gschwend et al., 2015; Shimshek et al., 2005). By contrast, the balance of redundancy and uniqueness is strongly shaped by mechanisms of presynaptic gain control, notably those mediated by periglomerular interneurons (Root et al., 2008; Vaaga et al., 2017; Zhou et al., 2020). These cells provide glomerulus-specific presynaptic inhibition, tuning the strength of the OSN-to-mitral/tufted cell synapse and adjusting the dynamic range of glomerular output. Importantly, parvalbumin-expressing interneurons (PV cells) have been shown to linearly transform mitral cell responses without strongly modulating their odour-tuning properties (Kato et al., 2013). Furthermore, neuromodulatory inputs, such as dopaminergic and GABAergic cotransmission, are known to alter gain and thus the spectrum of glomerular variance and decorrelation (Egger & Kuner, 2021; Vaaga et al., 2017). Manipulations targeting these gain control processes, either via pharmacological or genetic strategies, would be expected to systematically reshape the diagonal spectrum and provide direct tests of our model’s predictions.

In conclusion, by systematically comparing glomerular input and output responses across a large odour panel and linking these data to mechanistic models, we show that odour decorrelation in the mouse OB can be explained primarily by channel-specific feedforward scaling. This reframes a well-recognised transformation in the bulb as arising from simple gain control rather than requiring complex recurrent inhibition. These findings highlight a scalable rule by which early sensory circuits can enhance the separability of their inputs, while maintaining the flexibility to dynamically modulate representations through additional circuit mechanisms.

## Summary

We recorded glomerular input and output activity from dorsal OB glomeruli in response to a chemically diverse panel of ~50 odours. Output activity patterns were consistently less correlated than inputs and supported higher decoding accuracy, with the performance gain directly attributable to decorrelation. To probe the circuit basis of this transformation, we used a linear model of the OB constrained by our input data. Surprisingly, diagonal connectivity motifs, corresponding to channel-specific gain modulation without lateral connections, were nearly as effective as fully unconstrained models in reproducing the observed input–output transformation. Allowing lateral connectivity improved performance only modestly and did so through a small number of low-rank modes. These results indicate that feedforward gain modulation is sufficient to account for most of the decorrelation observed experimentally. Our work thus identifies feedforward scaling as an important computational operation in the OB, offering a simple yet powerful mechanism for enhancing odour separability and supporting ethologically relevant computations.

## Supporting information

Supplementary figures (high res)

## Methods

### Animals

Animals in this study were all 8–13-week-old mice (anaesthetised) and 14–22-week-old mice (awake) of C57/Bl6 background and mixed gender. For imaging sensory input, we used transgenic mice resulting of either MOR174/9-eGFP87 (Sosulski et al., 2011) or M72-IRES-EGFP (Potter et al., 2001) (JAX stock #007766) crossed into a Tbet-cre driver line (Haddad et al., 2013) (JAX stock #024507) crossed with a GCaMP6f reporter line (Madisen, 2015) (JAX stock #028865). For recording sensory output, we used transgenic mice resulting of M72-IRES-ChR2-YFP (Smear et al., 2013) (JAX stock #021206) or MOR174/9-eGFP87 crossed into a Tbet-cre driver line (Haddad, 2013) (JAX stock #024507) crossed with a GCaMP6f reporter line (Madisen et al., 2015) (JAX stock #028865).

Glomerular sensory input signals were recorded from a total of 5 mice (3 OMP-cre:MOR174/9-eGFP87-EGFP:Ai95 mice, 2 OMP-cre:M72-ChR2-YFP:Ai95 mice) and output signals from 12 mice (Tbet-cre:M72-ChR2-YFP:Ai95).

All animal protocols were approved by the Ethics Committee of the board of the Francis Crick Institute and the United Kingdom Home Office under the Animals (Scientific Procedures) Act 1986.

### In vivo imaging

#### Surgical and experimental procedures

Prior to surgery all utilised surfaces and apparatus were sterilised with 1% trigene. Mice were anaesthetised using a mixture of fentanyl/midazolam/medetomidine (0.05, 5, 0.5 mg/kg respectively). Depth of anaesthesia was monitored throughout the procedure by testing the toe-pinch reflex. The fur over the skull and at the base of the neck was shaved away and the skin cleaned with 1% chlorhexidine scrub. Mice were then placed on a thermoregulator (DC Temperature Controller, FHC, ME USA) heat pad controlled by a temperature probe inserted rectally. While on the heat pad, the head of the animal was held in place with a set of ear bars. The scalp was incised and pulled away from the skull with four arterial clamps at each corner of the incision. A custom head-fixation implant was attached to the base of the skull with medical super glue (Vetbond, 3M, Maplewood MN, USA) such that its most anterior point rested approximately 0.5 mm posterior to the bregma line. Dental cement (Paladur, Heraeus Kulzer GmbH, Hanau, Germany; Simplex Rapid Liquid, Associated Dental Products Ltd., Swindon, UK) was then applied around the edges of the implant to ensure firm adhesion to the skull. A craniotomy over the left olfactory bulb (approximately 2 × 2 mm) was made with a dental drill (Success 40, Osada, Tokyo, Japan) and then immersed in ACSF (NaCl (125 mM), KCl (5 mM), HEPES (10 mM), pH adjusted to 7.4 with NaOH, MgSO4.7H2O (2 mM), CaCl2.2H2O (2 mM), glucose (10 mM)) before removing the skull with forceps. The dura was then peeled back using fine forceps. A layer of 2% low-melt agarose diluted in ACSF was applied over the exposed brain surface before placing a glass window cut from a cover slip (borosilicate glass #1 thickness [150 µm]) using a diamond scalpel (Sigma-Aldrich) over the craniotomy. The edges of the window were then glued with medical super glue (Vetbond, 3M, Maplewood MN, USA) to the skull. Following surgery, mice were placed in a custom head-fixation apparatus and transferred to a two-photon microscope rig along with the heat pad. The microscope (Scientifica Multiphoton VivoScope) was coupled with a MaiTai DeepSee laser (Spectra Physics, Santa Clara, CA) tuned to 940 nm (<30 mW average power on the sample) for imaging. Images (512 × 512 pixels, field of view 550 × 550 µm) were acquired with a resonant scanner at a frame rate of 30 Hz using a 16 × 0.8 NA water-immersion objective (Nikon). Using a piezo motor (PI Instruments, UK) connected to the objective, a volume of ~300 µm was divided into 4 (sensory input imaging) or 12 planes (output imaging) resulting in an effective volume repetition rate of ~7.75 and ~2.5 Hz, respectively. The odour port was adjusted to approximately 1 cm away from the ipsilateral nostril to the imaging window, and a flow sensor (A3100, Honeywell, NC, USA) was placed to the contralateral nostril for continuous respiration recording and digitised with a Power 1401 ADC board (CED, Cambridge, UK).

For awake imaging experiments, mice were anaesthetised with inhaled isoflurane (1.5–2%). A bilateral craniotomy (2.5 × 2.5 mm over each hemisphere) was performed, although imaging was restricted to the left olfactory bulb. No agarose was applied prior to window implantation. Following surgery, animals were allowed to recover for at least 10 days before imaging sessions.

#### Odour stimulation

Odour stimuli were delivered using a set of custom-made 6-channel airflow dilution olfactometers. In brief, volumes of 3 ml of a set of 47 monomolecular odours (Sigma-Aldrich, St. Louis MO, USA) were pipetted freshly for each experimental day into 15 ml glass vials (27160-U, Sigma-Aldrich, St. Louis MO, USA). Odours were diluted 1:20 in air before being presented to the animal at 0.3 litres/min using custom Python Software (PulseBoy; github.com/RoboDoig). Odours were prepared for 3 s in tubing before a final odour valve was triggered to open at the beginning of an inhalation cycle for 2 s. During anaesthetised recordings all stimuli were presented with a 30 s inter-stimulus interval. During awake recordings, odour presentation lasted for 500 ms with a ~10s inter-stimulus interval. To minimise contamination between odour presentations, a high flow clean air stream was passed through the olfactometer manifolds during this time. Before each experiment, stimuli were calibrated using a photoionization detector (miniPID, Aurora Scientific, Aurora, ON, Canada) so that the concentration profile closely followed a final valve opening.

#### Data preprocessing

Motion correction, segmentation and trace extraction were performed using the Suite2p package (https://github.com/MouseLand/suite2p). Regions of Interest (ROIs) corresponding to glomeruli were manually delineated based on the mean fluorescence image in Fiji. The fluorescence signal from all pixels within each ROI was averaged and extracted as a time series with ΔF/F= (F - F0)/F0, where F = raw fluorescence and F0 = median of the fluorescence signal distribution. A custom MATLAB script identified duplicate glomeruli recorded in multiple imaging planes as those whose odour responses, computed as the integral of the change of fluorescence over the 5 s window following odour onset, had a Pearson correlation above 0.9, and that were colocalized within 20 µm in both X and Y directions. The most superficial ROI among the duplicates was kept and the remainder discarded from further analysis. This procedure yielded 148 input glomeruli and 167 output glomeruli from 17 mice (18.5 ± 8.3 (mean ± s.d.; range 6-36) glomeruli per mouse).

To allow pooling of data across multiple experiments, we next standardised the ROI odour responses. We first trial-averaged the ROI calcium traces to each odour. Next, to normalise these calcium traces, we calculated their means and standard deviations over the baseline period, defined as the three seconds before odour onset. We subtracted these baseline means from the trial-averaged responses, and divided the result by the baseline standard deviations, arriving at z-scored traces such as those shown in **Figure 2**. To reduce these z-scored time courses to a single odour response, we summed the responses over the 5 seconds following odour onset. This yielded a single z-scored response for each ROI to each odour. We used these data to compute the input and output correlations, and as input to the model.

### Computational methods

#### Correlation matrices

To compute pattern correlation matrices, ROI responses were first averaged over trials, then standardised to each have zero mean and unit standard deviation when computed across odours. Population vectors were formed by concatenating the processed responses of all ROIs to a given odour. Pattern correlation matrices were built by computing the Pearson correlation between population vectors for each pair of odours. The dimensions of the resulting matrices were thus *n* x *n*, where *n* is the number of odours, and each matrix element reported the Pearson correlation of ROI responses to the corresponding odour pair. To aid visual detection of patterns in the resulting matrices, the display sequence of odours was determined by hierarchical clustering based on the Euclidean distance of the input or output data.

#### Response variance

To quantify the overall response variance at the input and output levels, we computed the variance of each glomerulus’s response across the full odour panel (see **Figure 3D**). Specifically, we used the glomerulus-by-odour response matrix and obtained the covariance matrix across odours. The diagonal elements of this covariance matrix correspond to the variance of individual glomeruli across all odours. We then averaged these values across all recorded glomeruli to yield the overall response variance.

#### Classifier analysis

The data was split into a training (60%) and a test set (40%) using the StratifiedShuffleSplit function of the Python scikit-learn library. We trained linear Support Vector Machine (SVM) classifiers using an L1 penalty to promote sparse solutions, with a squared hinge loss function. The penalty parameter was fixed at C = 0.01. Classifiers were tested on the remaining 40% of trials. The trials to be saved for testing were picked at random. This training and testing were repeated 20 times with a random selection of testing trials used each time. The average performance over these 20 trials yielded the classification accuracy. To determine chance accuracy, classifiers were trained on the same data but with labels shuffled. As there were 48 stimuli (47 odours and mineral oil as blank), chance accuracy was at 1/48 ≈ 2.083%. To test how accuracy varied with the number of ROIs, we used the exact same procedure but used random subsets of ROIs of increasing number.

#### Principal component analysis

We performed principal component analysis (PCA) on odour responses of individual trials for the input and output data. Where necessary, we equalised the number of trials to 5 repetitions for all ROIs by adding up to 2 trials calculated from the median response of the individual ROI to an individual odour.

#### Transforming datasets

To transform the input dataset to the output, we first randomly subsampled ROIs and trials from the output to match the number of ROIs and trials on the input. This yielded two datasets containing the responses of 148 ROIs to 3 trials of 47 odours and mineral oil. Next, we computed the trial-average mean responses to each odour, and the scatter of the three trials around these means. We then sequentially transformed one dataset into the other, as described below. Three of the transformation steps involved changing the mean odour responses, the fourth, ‘scatter’, involved changing the single trial deviations relative to the mean responses.

##### Matching correlations and lengths

To transform one 148 ROIs x 48 stimuli matrix of mean odour responses, X, to match the lengths or correlations of another, Y, we first computed the Gram matrix Y’Y, of the latter. We then modified the elements of this Gram matrix based on whether we wanted to match length or correlations (described below) to yield a target Gram matrix G. Then we searched for the matrix Z closest in Frobenius norm to X but with the desired Gram matrix. That is, we solved the following problem

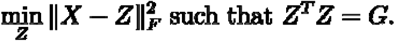

The solution to this problem is 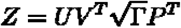, where ***U*** and ***V*** are the left and right singular vectors of ***X***, and ***P*** and **Γ** are the eigenvectors and (diagonalized) eigenvalues, respectively, of ***G***.

To determine the Gram matrices for matching correlations or lengths, we first decomposed G as the elementwise (Hadamard) product of a matrix of cosine correlations and a rank-1 matrix of pairwise length products. That is,

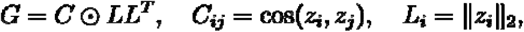

where ***z***_***i***_ is the mean odour response to the i’th odour after the transformation. To find the nearest matrix to X that had the same lengths as Y while preserving the correlations in X, we set ***C***_***ij***_ = **cos** (***x***_***i***_, ***x***_***j***_), and ***L***_***i***_ = **‖ *y***_***i***_ **‖** _**2**_, where ***x***_***i***_ is the response to the i’th odour in the dataset X, and similarly for ***y***_***i***_. To find the nearest matrix to X that had the same correlations as Y but maintained the lengths of X, we set ***C***_***ij***_ = **cos** (***y***_***i***_, ***y***_***j***_), and ***L***_***i***_ = **‖ *x***_***i***_ **‖** _**2**_.

##### Matching rotations

Rotating a matrix of means X to match a second matrix of means Y corresponds to solving the problem

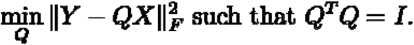

This problem has the solution ***Q* = *VU***^***T***^, where ***U*** and ***V*** are the left and right singular vectors of ***XY***^***T***^, respectively.

##### Matching scatter

To match the scatter of trials around their odour specific means we simply computed the deviation of each trial from its mean in one dataset, and added this deviation to the corresponding mean in the other dataset, to form the corresponding trial of data.

#### Modelling the Input–Output Transformation

##### Linear model formulation

To find connectivity that can produce the input–output transformation we observe, we considered a linear model of the olfactory bulb in which mitral cells were excited by their glomerular inputs and inhibited by one another. These dynamics can be represented compactly in matrix form as

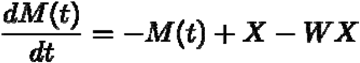

Here 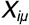 is the input to glomerulus ***i*** for odour ***μ*** and is assumed constant. 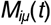 is the corresponding mitral cell activity at time ***t***. The strengths of this inhibition were set by the matrix ***W***, where ***W***_***ij***_ is the strength of the functional inhibition of mitral cell ***i*** by mitral cell ***j***.

To find the steady-state response of the mitral cells to the full bank of odours we set 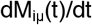 above to zero, which yields ***M*** = **(*I* + *W*)** ^**−1**^ ***X***.

This equation relates how inputs *X* to a set of glomeruli are transformed by the recurrent connectivity ***W*** into outputs *Y* of the same set of glomeruli. The expression above shows that the recurrent connectivity defines an equivalent feedforward connectivity, *Z*, defined as ***Z* ≜ (*I* + *W*)** ^**−1**^, which transforms the input to the output according to *M = ZX*. In what follows we will frequently use this equivalent feedforward model of the input–output transformation.

##### Fitting Representations

Given our model’s linear relationship of inputs to outputs, we would ordinarily now substitute our recorded inputs for *X*, and find the connectivity *Z* that produced outputs *M* closest to those, *Y*, that we actually recorded. However, we did not record glomerular inputs and outputs simultaneously, and the identity and number of glomeruli that we recorded from are different on the input and output sides. Therefore, rather than matching inputs and outputs at the level of individual ROIs, we instead focused on the pattern of population responses. Specifically, we aimed to reproduce the pattern of pairwise odour similarities, i.e. the *representation*, that we observed in the output population.

The similarity of odour responses can be measured in many ways. Because the key experimental observation we aimed to understand is the reduction in Pearson correlation from input to output, the natural measure of similarity would have been Pearson correlations. It is numerically straightforward to optimize Z to match representations using this measure of similarity. However, investigating the connectivity solutions analytically would be difficult. This is because computing Pearson correlations requires normalization of covariances by standard deviation, which introduces a nonlinearity that complicates the analysis.

To balance analytic tractability with relevance to our experimental observations, we chose to measure similarity by covariance alone. Because Pearson correlations can be computed from covariances, matching the latter exactly would automatically match the former as well. Therefore, we hoped that our compromise would yield analytically interpretable insight into the decorrelation we observed.

Our procedure therefore transformed input responses *X* into model outputs *ZX*, compared the resulting covariances to those for the output responses *Y*, and searched for the feedforward connectivity that minimized the sum of squares difference between the resulting odours-by-odour matrices. We expressed this search minimization of the following loss function,

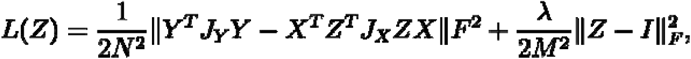

where *N* is the number of odours, *M* is the number of input (and hence model) glomeruli, ||.||^2^_*F*_ returns the sum of squares of the elements of its matrix argument, λ is a regularization parameter, and the matrices *J*_*Y*_ and *J*_*X*_ perform the mean-subtractions required when computing covariances.

The first term measures the difference between the predicted and observed output covariances. The second term is a regularizer that favours the default transformation in which the input is passed directly to the output, unchanged. This can be expressed in terms of feedforward connectivity as a bias towards the identity, as we do above. It can be equivalently expressed in terms of recurrent connectivity as a bias towards lack of recurrence i.e. W = 0.

##### Computing the Fits

To compute a connectivity solution, we started with the z-scored responses and first generated training, test and validation datasets. To do so, we noted that the number of repetitions of an odour varied across experiments, but that at least three were always available. Therefore, for either the input or output recordings, we generated a training set by sampling one trial at random per ROI and odour, a validation set by repeating the procedure on the remaining trials, and a test set by repeating the procedure once more on the rest of the trials.

For each of the trials generated above, we performed an additional normalization step. We normalized the output responses by their overall standard deviation to remove a scaling degree of freedom. We normalized the input responses to have unit variance per ROI. This normalization of inputs improved their trial-to-trial stability of their odour representations as measured by Pearson correlation. Because the un-normalized input response amplitudes varied significantly across ROIs, this normalization step helped avoid decorrelation solutions that merely equalized overall amplitudes in favour of those that were sensitive to the pattern of odour responses of each ROI.

We generated multiple datasets in this way, and computed fits (described below) for each of them, for each value of the relevant regularization parameter(s). We determined the best value of the regularization parameter as the one that yielded the lowest average error over our validation datasets. Finally, we applied the resulting optimal value of the regularization parameter to transform the inputs in the test set to outputs and report performance.

Given a set of input and output data, we minimize the loss function using ‘scipy.minimize’ with the truncated conjugate gradient (TNC) algorithm, parametrizing the loss function to reflect the structural constraints we impose. For example, when searching for connectivity constrained to lack lateral connections, we parameterize the loss function to receive the diagonal elements of the full connectivity matrix *Z*, setting the rest to zero. To speed convergence, we also provide scipy.minimize with manually-derived gradients of each parameterization of the loss function.

##### Diagonal-only model

In the diagonal-only model, lateral connections were excluded, so each channel could only be amplified or attenuated by a self-gain. To determine the logic of the gains is simple in principle - find the values of the gains for which the loss has gradient 0. However, this is complicated for our loss function: covariances are quadratic in the connectivity, therefore their squared differences are quartic, and gradients are one degree lower, cubic, in the connectivity parameters, making the results hard to interpret.

However, we found empirically that for the best cross-validated fits we needed large regularization. This in turn meant that our learned connectivity stayed near its regularization target, which in the case of diagonal connectivity was unit gain. We leveraged this observation and linearized our loss function around this value. This yielded an interpretable, closed form approximation for the gains, which we now describe.

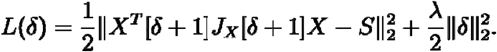

We first expressed our loss function in terms of the deviations from unit gain ***δ* = *z* − 1**, Here the square brackets are shorthand for a diagonal matrix formed of their arguments, and we’ve absorbed the various constants of proportionality in the original loss into the regularization parameter ***λ***. We have also rotated the odour space coordinates in the goodness of fit terms so to a basis in which the observed output covariance is diagonal, with variances given by the elements of the diagonal matrix ***S***.

We first consider the terms bracketing ***J***_***X***_ and keep only the contributions linear in ***δ***,

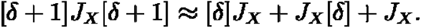

Now bringing in the interactions with ***X***,

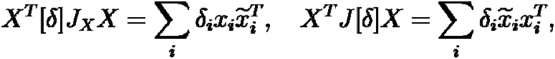

where 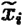 are the population-mean-subtracted responses.

We can sum the two terms above into a single linear expression in ***δ***_***i***_,

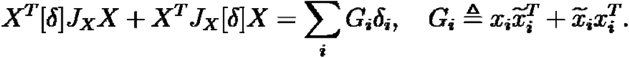

We can combine the last term with the target covariance as

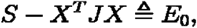

where the subscript reminds us that this is the error at ***δ* = 0**.

If we vectorize, forming

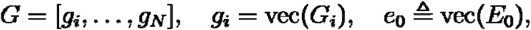

our loss will be

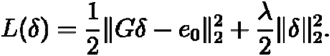

The gradient of this with respect to ***δ*** is

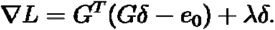

Setting this to 0 gives

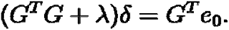

In terms of single units, this says

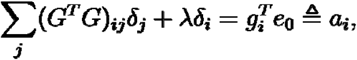

where ***a***_***i***_ is the unit’s alignment with residual ***e***_**0**_. Rearranging to isolate ***δ***_***i***_, we get

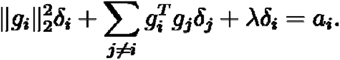

We can then solve for the gains as

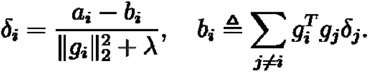

Therefore, the linearized gain applied to each unit is determined, first, how much each unit’s own representation ***g***_***i***_ overlaps with the difference in representations between the output and input, as measured in ***a***_***i***_. The second is how much the unit’s representation overlaps with that of the other units, weighted by the gains applied to those units, as measured in ***b***_***i***_.

##### Free connectivity

We also fit connectivity without any constraints beyond regularization. Expressed in the basis of the output covariance, this loss is

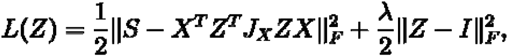

where ***S*** is the diagonal matrix of output variances, ***X*** is the input data rotated into the output odour space, and we’ve absorbed various constants into ***λ***.

The gradient of the loss with respect to ***Z*** is

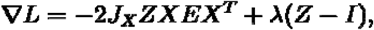

where we’ve defined the difference between the output covariance and the predicted covariance as

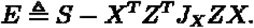

Note that this is symmetric, and quadratic in ***Z***.

Setting the gradient to zero characterizes the solution

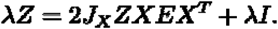

Notice that the right-hand side is cubic in ***Z***, since ***E*** is quadratic.

#### Symmetry

To show that the solutions we find are symmetric we will express ***Z*** in a basis that isolates the effect of the mean subtraction. In particular, we can decompose the identity as 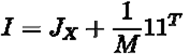. Notice that these two components are orthogonal. We can then decompose ***Z*** as

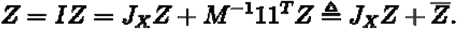

To find the first component at the solution, we multiply the solution condition on the left by ***J***_***X***_, and find

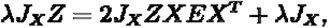

which then gives

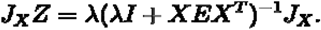

The columns of ***X*** are mean-subtracted, so we can decompose it using SVD as ***S*** = ***US***_***X***_***V***^***T***^ and orthonormally complete its column space ***U*** to span the full space of mean-subtracted vectors. In such a basis 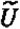, ***J***_***X***_, as the mean-subtraction operator, is the identity. The expression above in this basis is

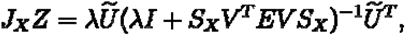

which is symmetric.

The remaining component of ***Z*** is 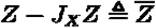, where the elements of each column have the same column-specific value, the mean of that column of ***Z***. The first term in the gradient is mean subtracted by ***J***_***X***_ so it doesn’t affect this component. The only contribution comes from the second term, so we get at the solution, 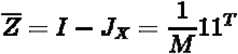 which is symmetric. Since 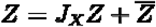, and both components are symmetric, ***Z*** itself is symmetric.

#### Approximating the solution

As we noted above, the equations that determine the free connectivity solution are cubic in ***Z***. Therefore, even if we could solve them in closed-form, the solutions would be complex and unlikely to yield insight. We therefore searched for an insightful, closed-form solution to the equations.

We again note that we needed high regularization for best cross-validation performance, and that our solution would likely remain close to the regularization target of the identity matrix. We therefore studied the solutions as deviations ***W*** from the identity.

Our initial approximation attempt was analogous to the diagonal connectivity approach: approximating the loss function to be quadratic in the deviations, yield linear approximate solutions. However, this did not give good results because the approach works best when the true deviations are small, which was the case for many, but not all elements of the true learned connectivity.

Therefore, we tried a different approach, in which we maintained the full, quartic loss, but approximated the solution along the regularization path. We first switched our parameterization of the loss to be in terms of the inverse regularization parameter, ***δ***^**−1**^ **≜ *ε***,

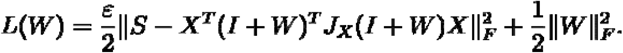

Setting the gradient of this loss to zero gives the full solution condition as

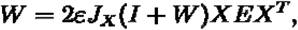

where ***E*** is defined as in the previous section. Note again that this is a cubic equation as ***E*** is quadratic in ***W***.

Each value of inverse regularization ***ε*** will give its own solution, if one exists. We can index these as ***W* (*ε*)**. In particular, ***W*(0)** corresponds to the solution when the regularization is infinitely strong, and the deviations are forced to be 0.

The solution path we describe above is for the exact solutions. To achieve our approximation, we noted that we needed large regularization, therefore small inverse regularization. We therefore approximated our exact solution as

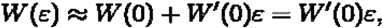

since as discussed above ***W*(0) =0**. Here ***W ′* (0)** is the derivative along the solution path, evaluated at ***ε* =0**. We compute it as

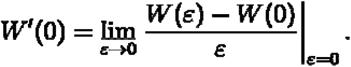

Using ***W*(0) =0** and substituting our expression above for ***W* (*ε*)** we get

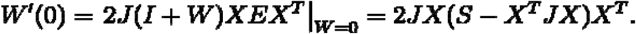

Since our input data are mean-subtracted, ***J***_***X***_***X =X***, and we arrive at our approximate solution

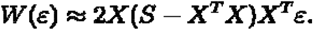

In **Figure 7** we compare the modes of this approximation to those of the full solution.

## Data availability

Source data supporting the graphs presented in the main figures is available at https://zenodo.org/records/17359151.

## Code availability

Analysis code to recreate the main figures is available from the following repositories:

https://github.com/ackels-lab/ob_gain_control_decorrelation, https://github.com/stootoon/ob_io_conn_models, and https://github.com/stootoon/glom-io-transform-release.

All custom analysis code supporting the findings of this study will be made publicly available through the same code repositories upon publication.

## Acknowledgements

We thank Yuxin Zhang for her help with data extraction and curation in the awake recordings. We also thank all members of our laboratories for helpful discussions and support throughout this work. This work was supported by the Francis Crick Institute which receives its core funding from Cancer Research UK (FC001153), the UK Medical Research Council (FC001153), and the Wellcome Trust (FC001153); by a Physics of Life grant (EP/W024292/1) funded by EPSRC and Wellcome; by the NSF/CIHR/DFG/FRQ/UKRI-MRC Next Generation Networks for Neuroscience (“NeuroNex”) Program “From Odor to Action” (Award #2014217); A.T.S.), a BIF doctoral fellowship to Y.Y., and a DFG postdoctoral fellowship to T.A. It was further supported by the German Research Foundation (FOR5424 “Modolfor”, A.T.S and T.A) and the European Union (ERC, “TempCOdE”, 101077017, T.A.). Views and opinions expressed are, however, those of the author(s) only and do not necessarily reflect those of the European Union or the European Research Council. Neither the European Union nor the granting authority can be held responsible for them.

## Supplementary figures

**Figure S1.1:**
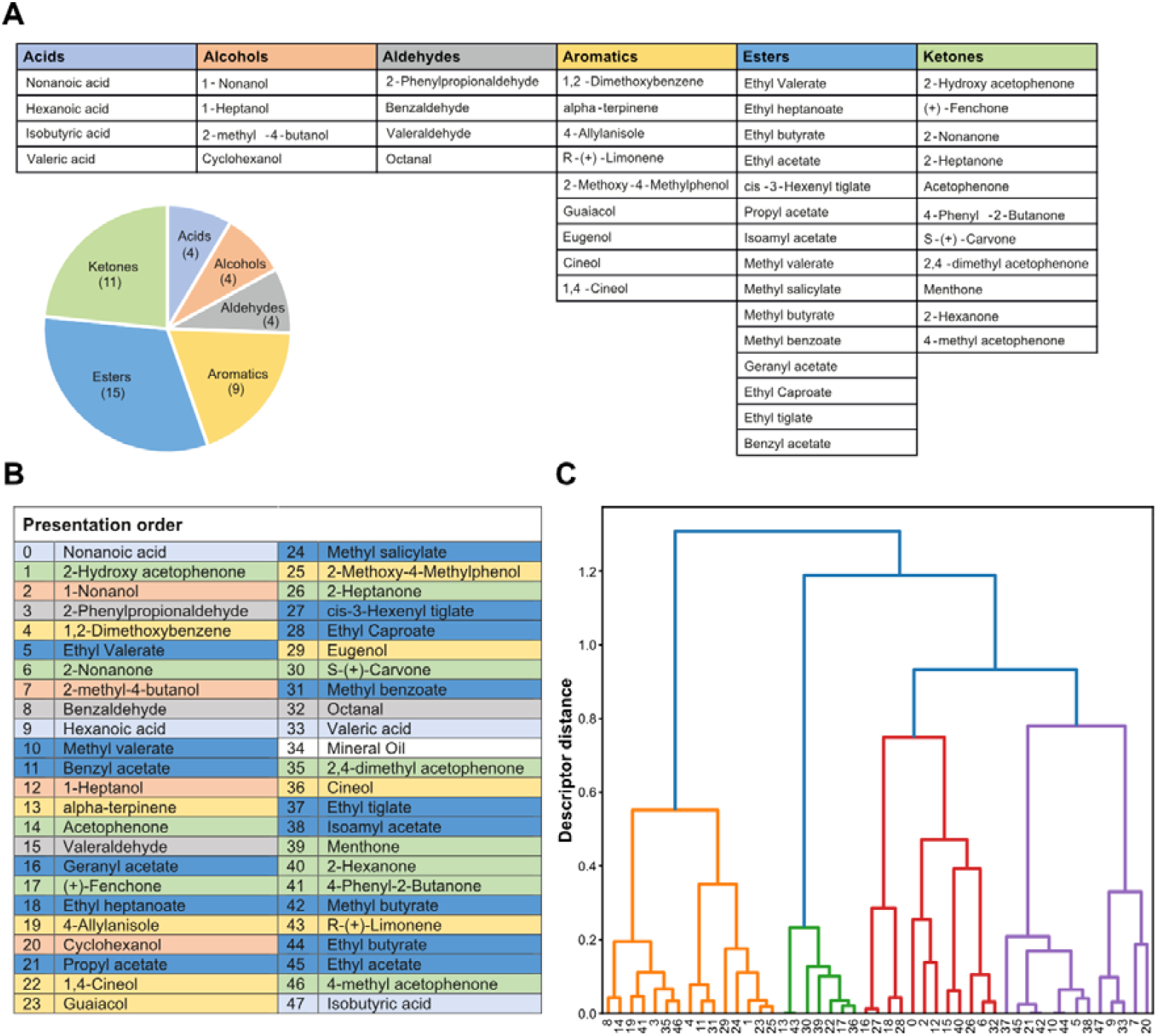
Experimental odour panel. **(A)** List of monomolecular odours presented during experiments sorted by chemical class. **(B)** Sequences of odours as presented during the experiment. **(C)** Molecular descriptor-based clustering of the odour panel. Hierarchical clustering dendrogram based on 9 molecular descriptors (correlation distance, average linkage). Colours indicate chemical class. Numbers show odour IDs.

**Figure S1.2:**
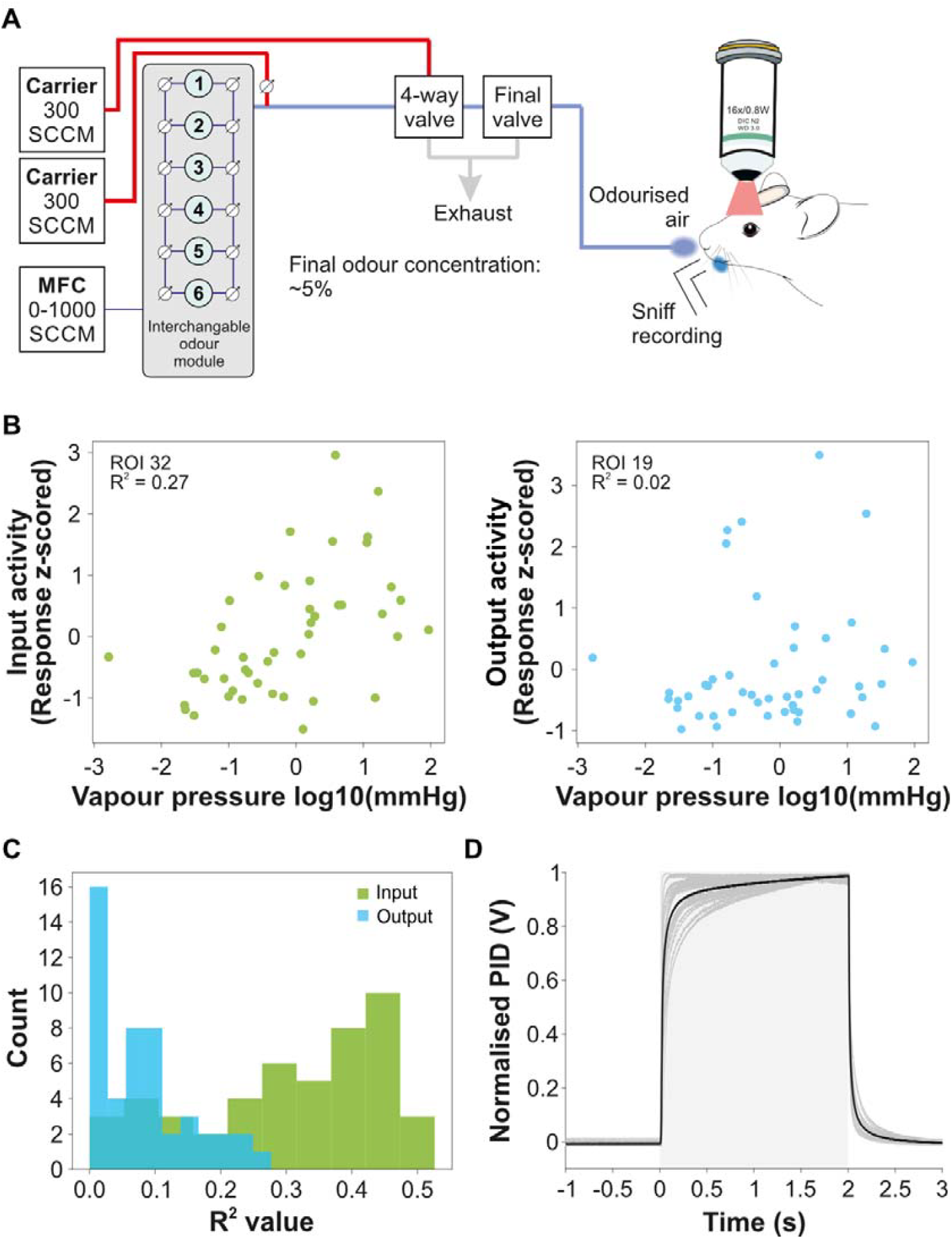
Schematic of odour presentation and stimulus characteristics. **(A)** Schematic of the custom odour delivery system. **(B)** Z-scored odour response of an example input (left) and output glomerulus (right) plotted as a function of odour vapour pressure shown in a log10 scale. **(C)** Histogram of R-squared values for all glomeruli computed by linearly regressing each glomerular z-scored response as a function of vapour pressure. **(D)** Overlay of temporal profiles of 21 odours as recorded using photoionisation detector (PID). Shown are recordings of all 21 monomolecular odours that elicited a voltage signal in the PID (grey traces) and the average signal of these odours (black trace). Odour presentation period from 0-2 seconds (grey shaded area), traces are normalised between 0 and 1.

**Figure S2.1:**
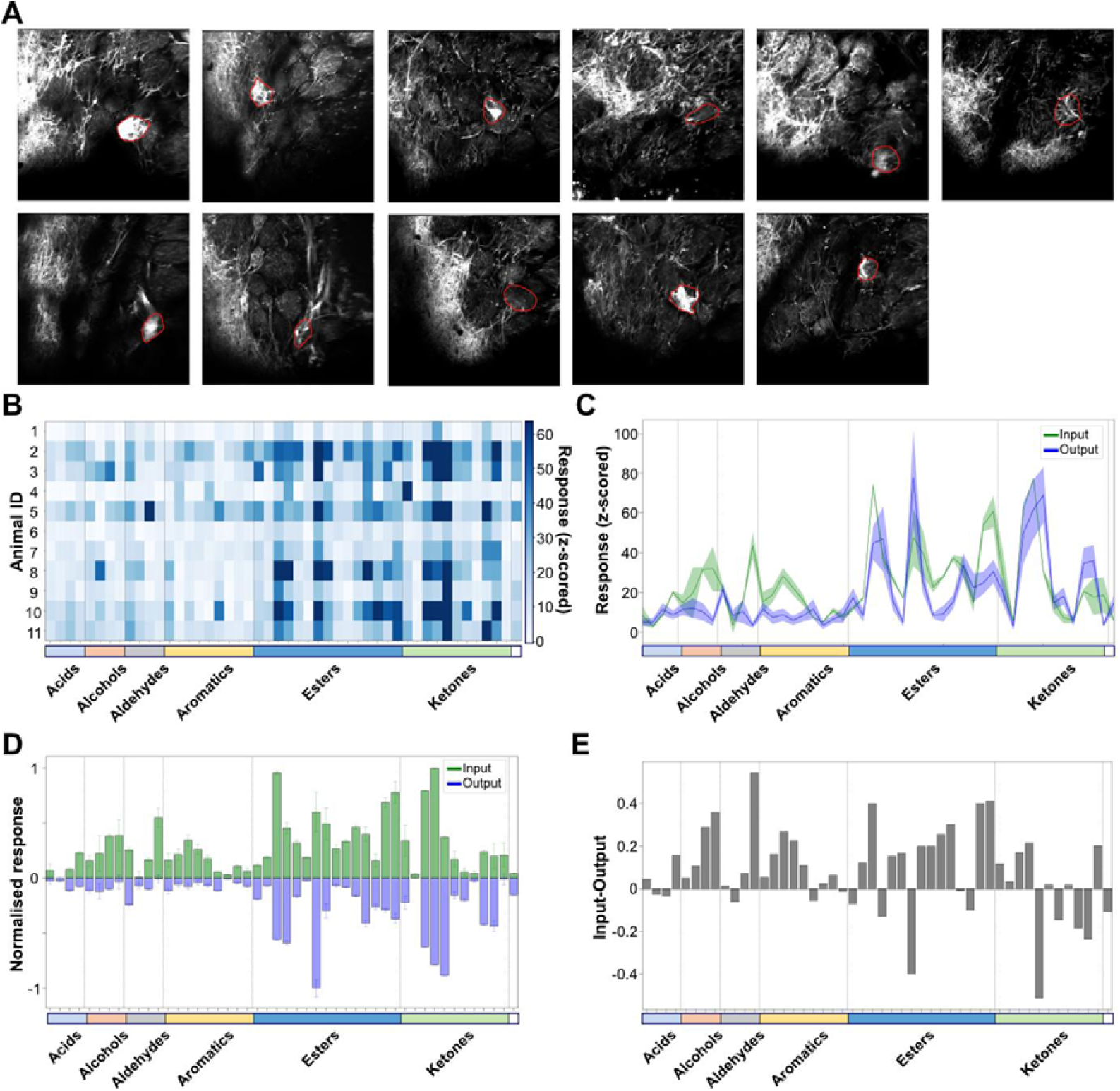
Input–output transformation of M72 glomerular odour responses. **(A)** Fluorescence from output neuron dendrites recorded in the glomerular layer of Tbet-cre:M72-ChR2-YFP:Ai95 mice. The M72 glomerulus is identified by YFP-labelled sensory axons (red contour; n = 11 animals). **(B)** Heatmap of individual M72 output responses (z-scored and integrated) to all odours sorted by chemical class. **(C)** Average z-scored responses to all odours as measured from the M72 glomerulus (mean ± SEM, input: green, output: blue). **(D)** Normalised average responses to all odours presented as mean ± s.d. (input: green, plotted as positive; output: blue, plotted as negative). **(E)** M72 response profile plotted as difference between input and output signal. Input: n = 2 animals, output: 11 animals.

**Figure S3.1:**
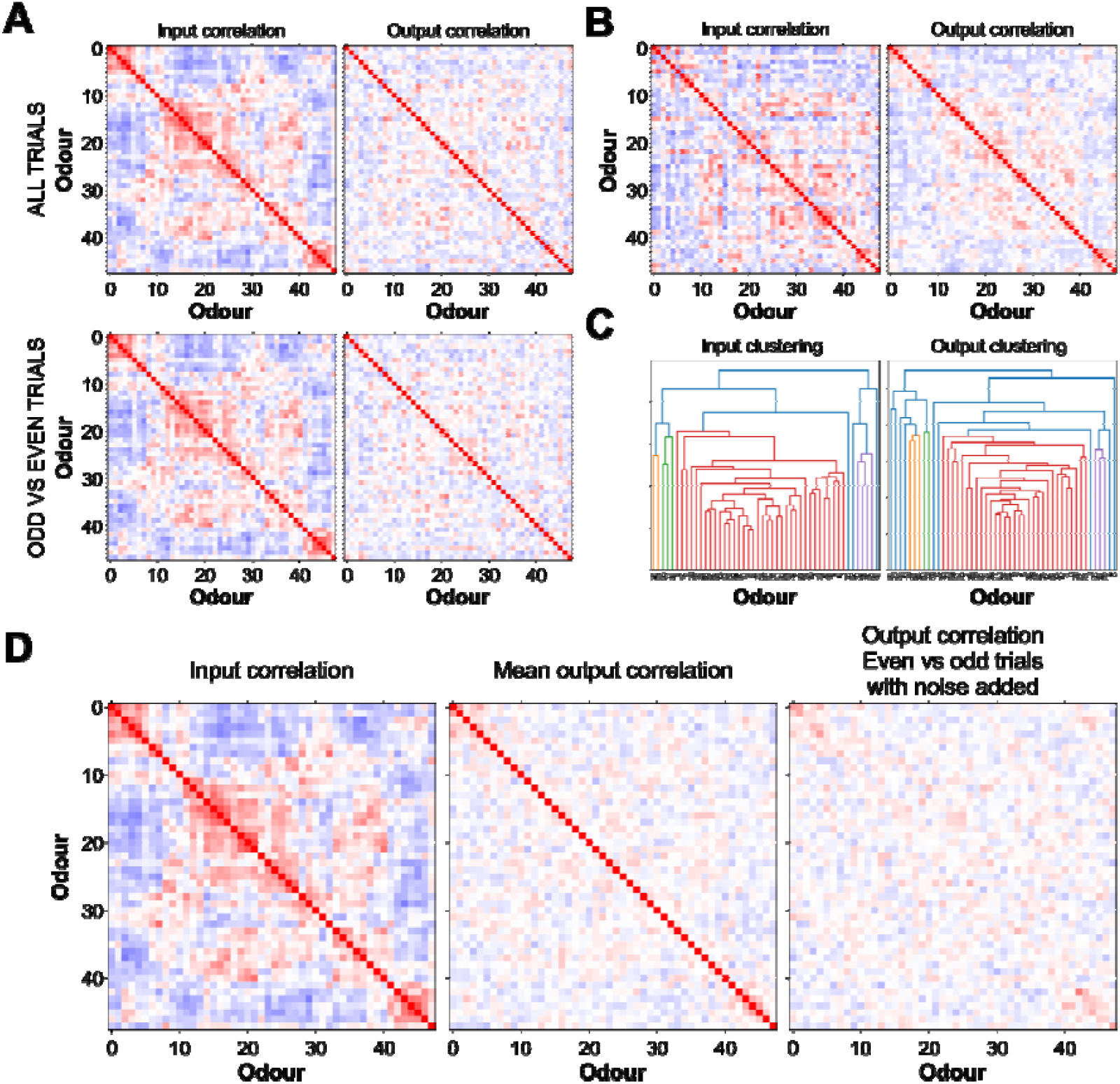
Decorrelation is consistent across trials and is not based on random noise. **(A)** Correlation matrices for input and output data computed by using all trials (top row) and by correlating odd against even trials. **(B)** Correlation matrices using odour sequence determined by hierarchical clustering based on the Euclidean distance of the output data. **(C)** Dendrograms resulting from hierarchical clustering of input (left) and output data (right). **(D)** The addition of random noise (2 s.d.) to the raw data before computing the correlation matrix of the output data reduces the cross correlation to approximately zero on the diagonal, proving that the decorrelation is not down to random noise.

**Figure S3.2:**
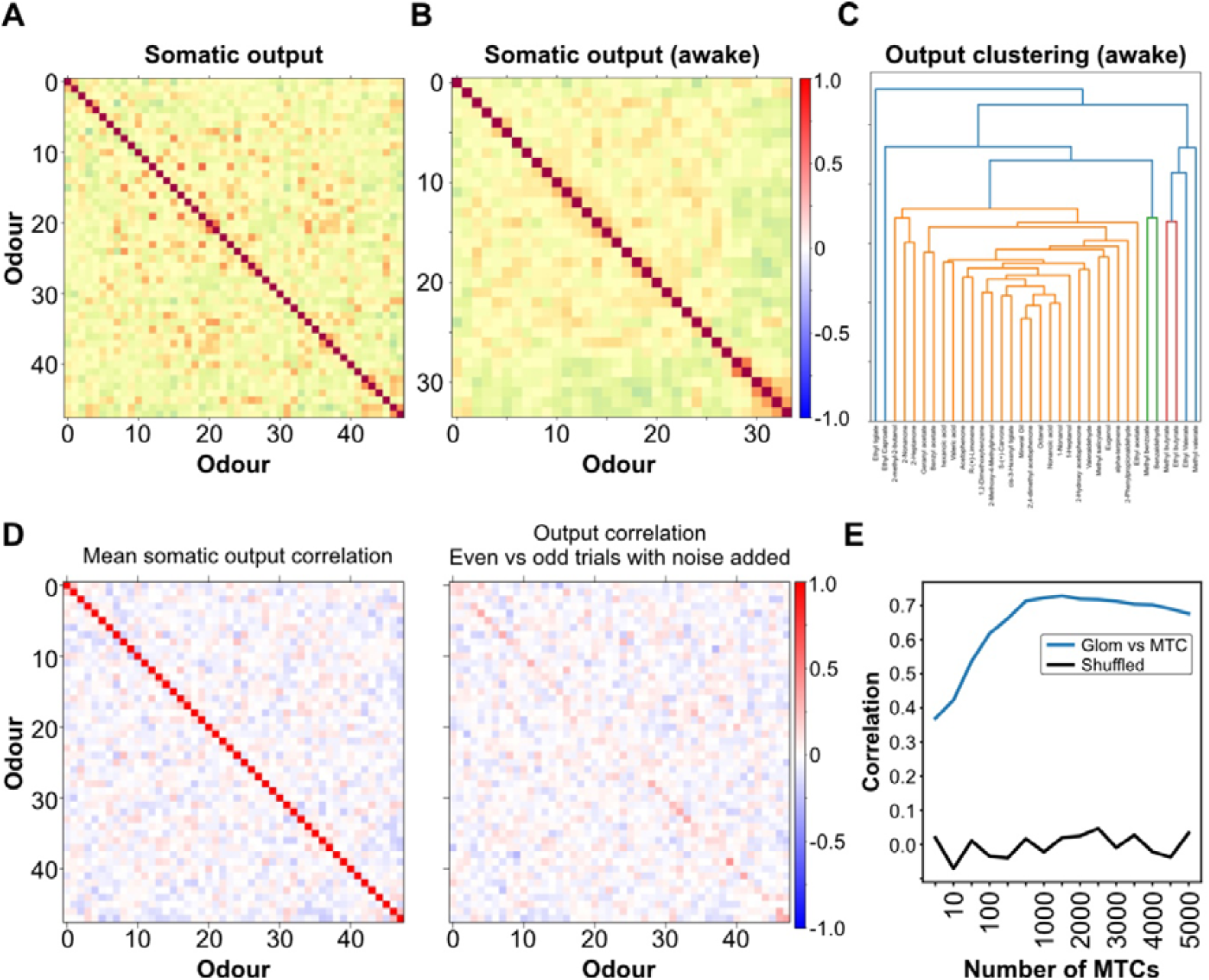
Pattern correlation of somatic odour responses. **(A)** Odour correlation maps of somatic ROIs. The sequence of odours was determined using hierarchical clustering based on the Euclidean distance of the input data. To equalise the number of ROIs for both datasets, 167 somatic ROIs that showed the highest variance across odour responses were selected. **(B)** Odour correlation (34 odours) for somatic ROIs (n = 167 MTCs) recorded in awake mice, rendering a comparable correlation structure to the anaesthetised state. **(C)** Hierarchical clustering used to define the sequence of odours shown in (B). **(D)** The addition of random noise (2 s.d.) to the correlation matrix of the output data reduces the cross correlation to approximately zero on the diagonal, proving that the decorrelation is not down to random noise. **(E)** Correlation between glomerular and somatic correlation matrices as a function of the number of M/T cells that were used. The correlation was calculated using the off-diagonal values of correlation matrices (blue: original order, black: order of correlation shuffled row wise).

**Figure S3.3:**
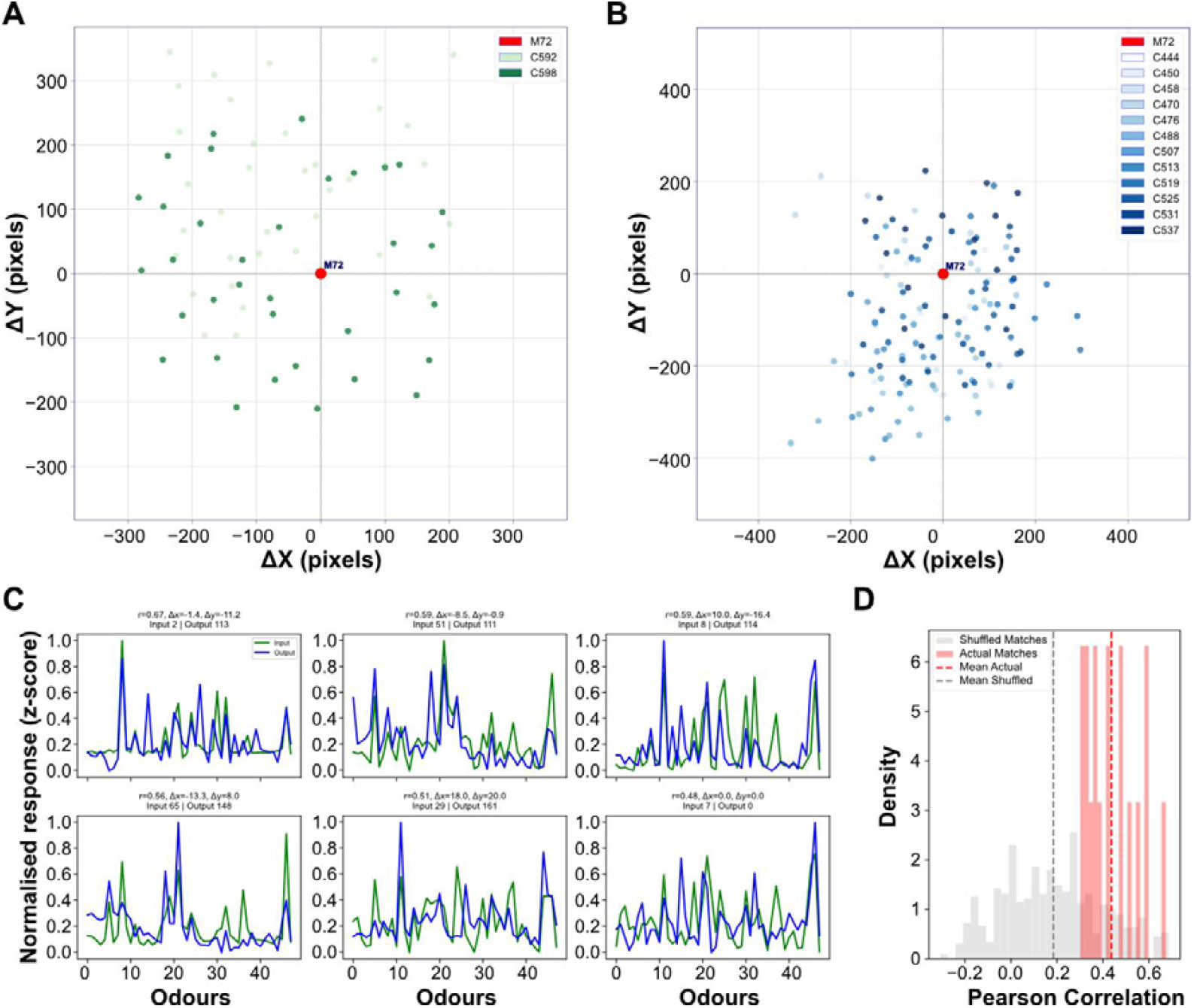
Spatial and functional matching of glomeruli centred around the M72 glomerulus. **(A)** Spatial distribution of glomerular input ROIs from two animals, aligned to the centroid of the M72 glomerulus (red). Colour intensity reflects different animals. **(B)** Same as in (A), but showing the distribution of output ROIs from twelve animals. **(C)** Odour response profiles of nine representative input–output ROI matched pairs with high correlation, sorted by descending Pearson correlation (r). Input and output responses are shown in green and blue, respectively. **(D)** Distribution of Pearson correlation coefficients for functionally matched ROI pairs with r > 0.3 (red) and shuffled controls (grey). Matched pairs showed significantly higher correlation than shuffled controls (Kolmogorov-Smirnov test, p = 1.73e-09; KS statistic = 0.721), suggesting consistent matching across input and output representations centred on M72.

**Figure S3.4:**
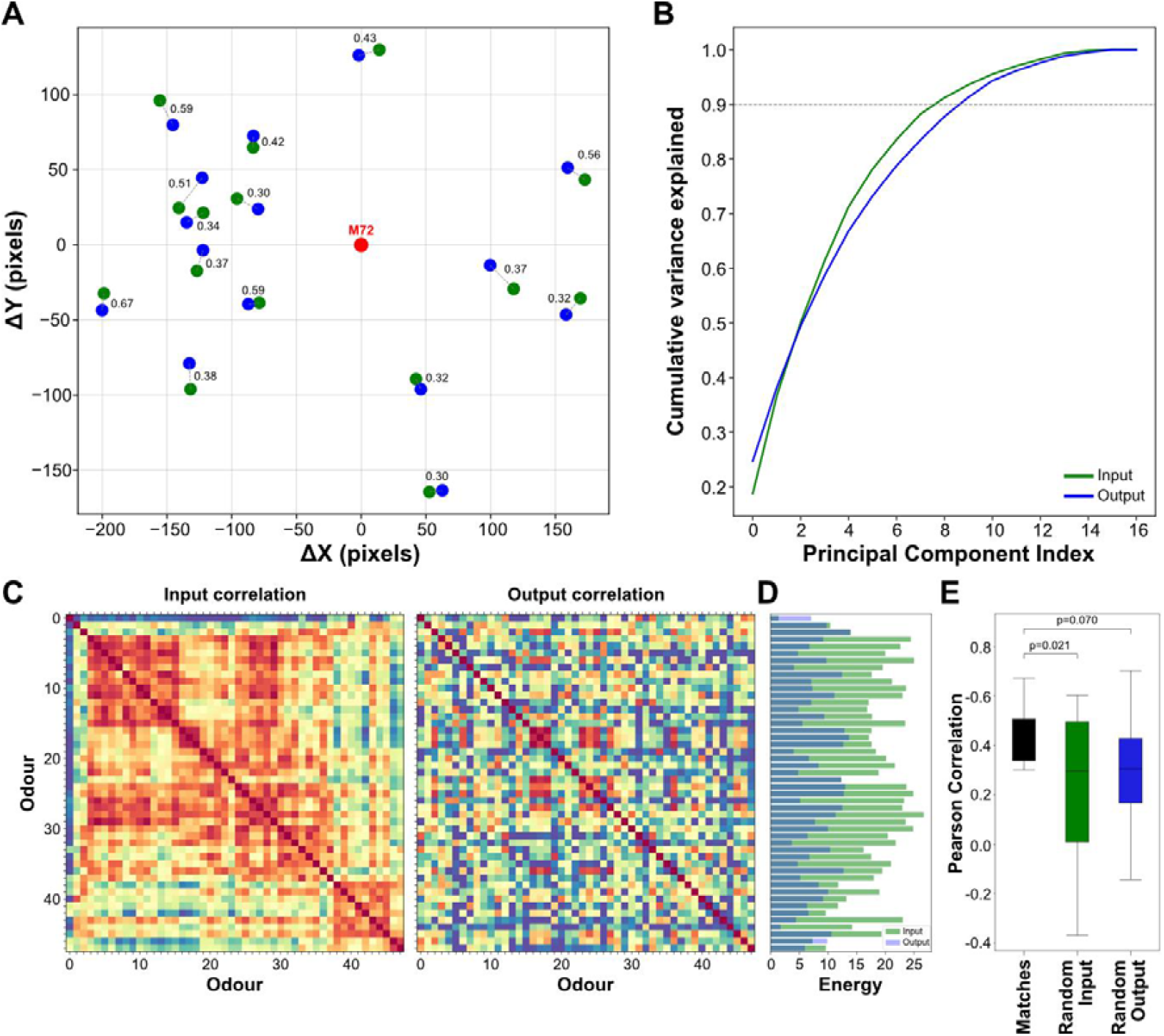
Odour representations of matched glomeruli become decorrelated from input to output. **(A)** Spatial distribution of input (green) and output (blue) ROIs matched by both spatial proximity and functional similarity (Pearson correlation r > 0.3), aligned to the M72 glomerulus (red) as a common reference point. Correlation values for each matched pair are indicated. **(B)** Cumulative variance explained by principal component analysis (PCA) of matched glomerular odour responses. Input responses (green) reached 90% variance with 9 components compared to output responses (blue, PC 10), indicating larger dimensionality in output representations for the subset of matched glomeruli. **(C)** Odour correlation matrices for matched input (left) and output (right) glomeruli (n = 17). Odour order was fixed across both matrices using hierarchical clustering based on the input data to facilitate direct comparison. The output matrix exhibits less structure and lower correlation, consistent with decorrelation. **(D)** Energy of odour correlations (defined as the squared sum of pairwise correlations minus 1) for each odour, comparing input (green) and output (blue). Most odours show reduced correlation energy in the output, further supporting a broad decorrelation of glomerular representations. **(E)** Correlation values between spatially matched glomeruli are significantly higher than correlations with randomly paired input or output ROIs (n = 17 ROIs).

**Figure S4.1:**
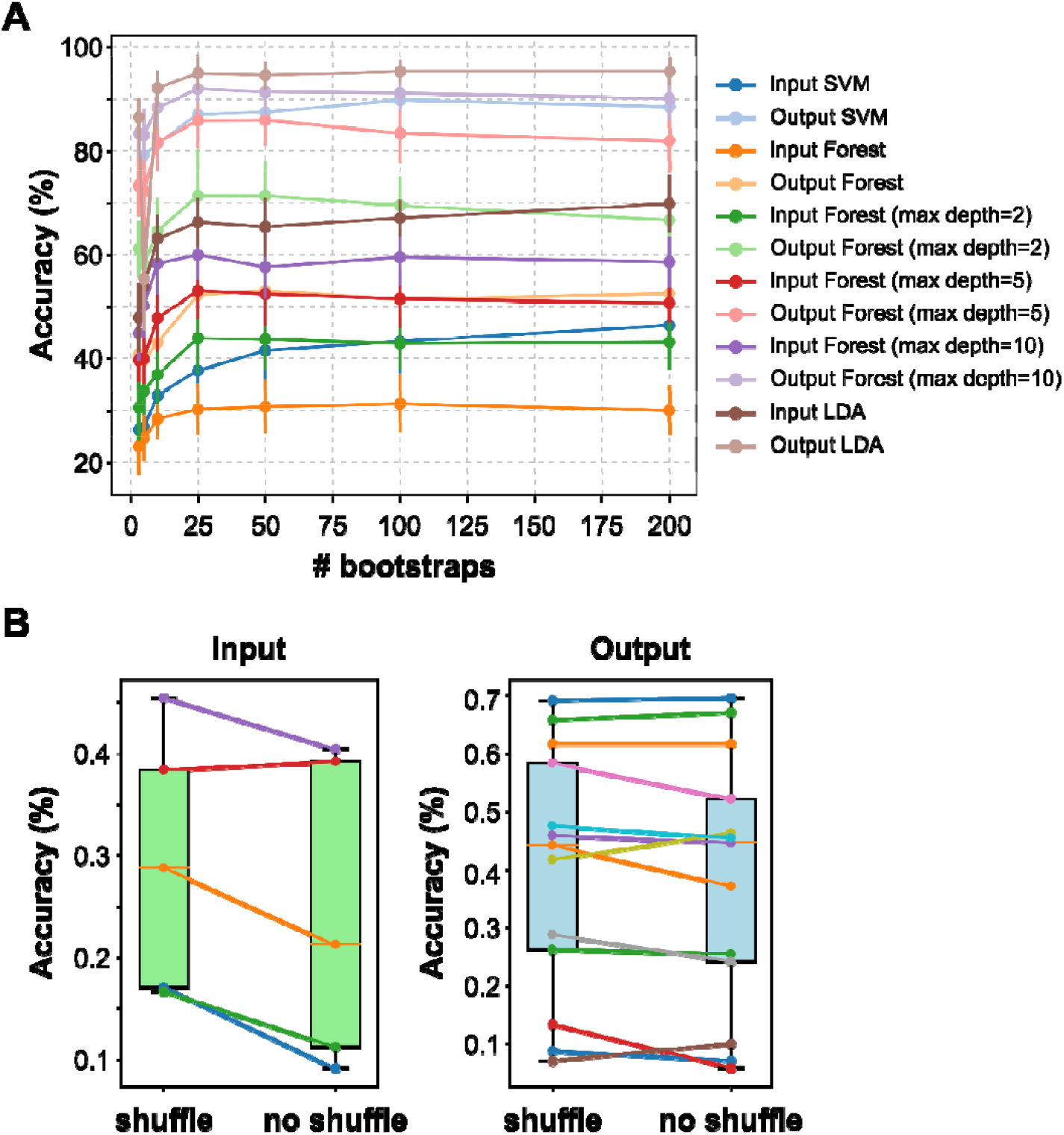
Effect of bootstrapping and label shuffling on classifier accuracy. **(A)** Comparison of different classifiers (SVM, Random Forest and LDA) on odour prediction accuracy as a function of the number of bootstrapped odour repetition trials. Accuracy plateaus for most classifiers at around 25 trials. **(B)** Shuffling trial order and thereby omitting noise correlations when using 3-5 experimentally acquired trials has no significant effect on prediction accuracy (input shuffle: 0.29 ± 0.11, input no shuffle: 0.24 ± 0.13, mean ± s.d., p = 0.54; output shuffle: 0.42 ± 0.21, output no shuffle: 0.39 ± 0.21, mean ± s.d., p = 0.78).

**Figure S5.1:**
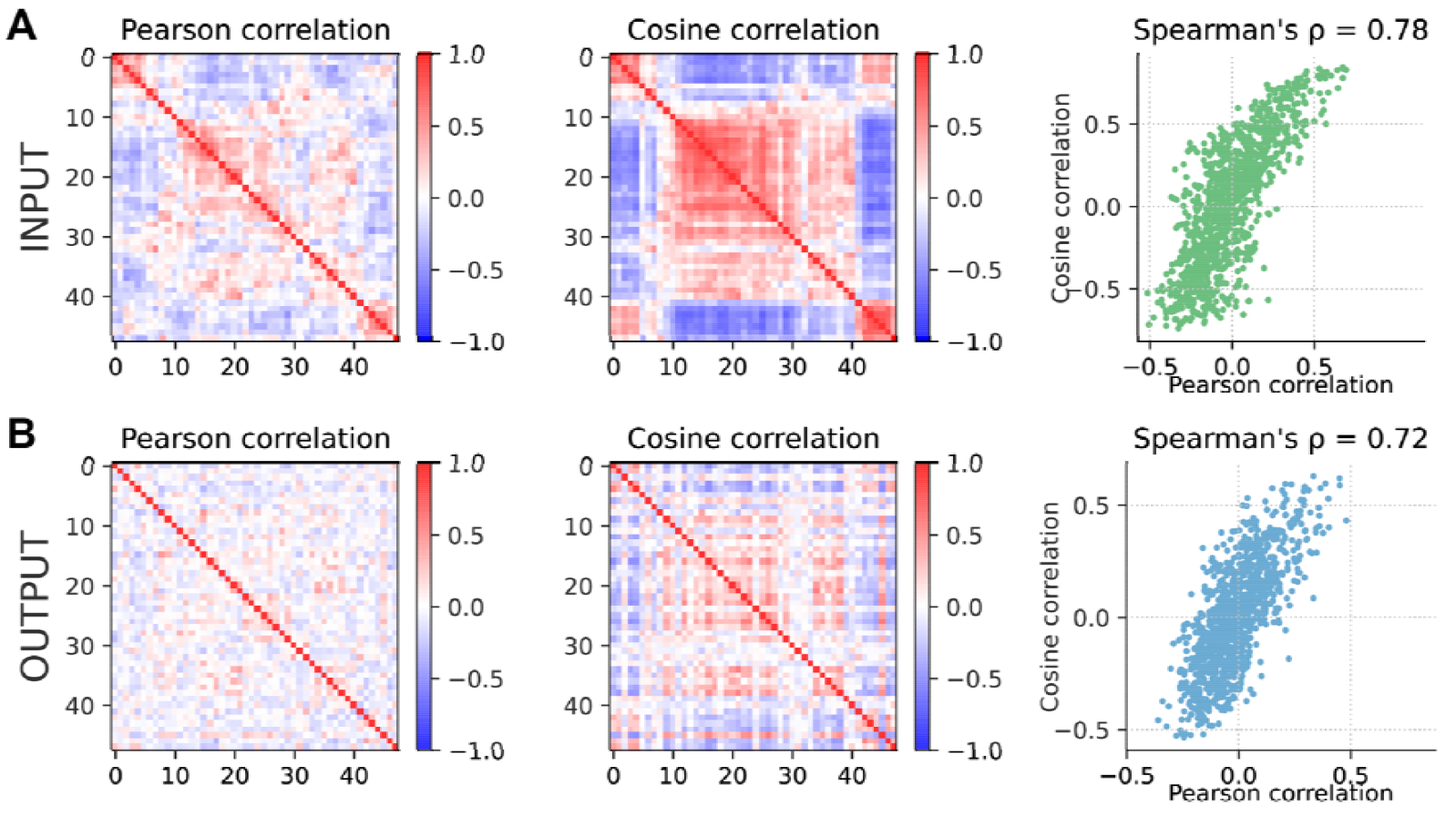
Pearson correlation and cosine correlation are correlated. **(A)** Pearson correlations (left) and cosine correlations (middle) of trial-averaged odour responses of olfactory bulb inputs. Ticks indicate odours, ordered as in the main text. Pearson correlation plots are as in the Main Text; right: Pearson correlation vs. cosine correlation for all pairs of odours. **(B)** As in (A) but for olfactory bulb outputs.

**Figure S5.2:**
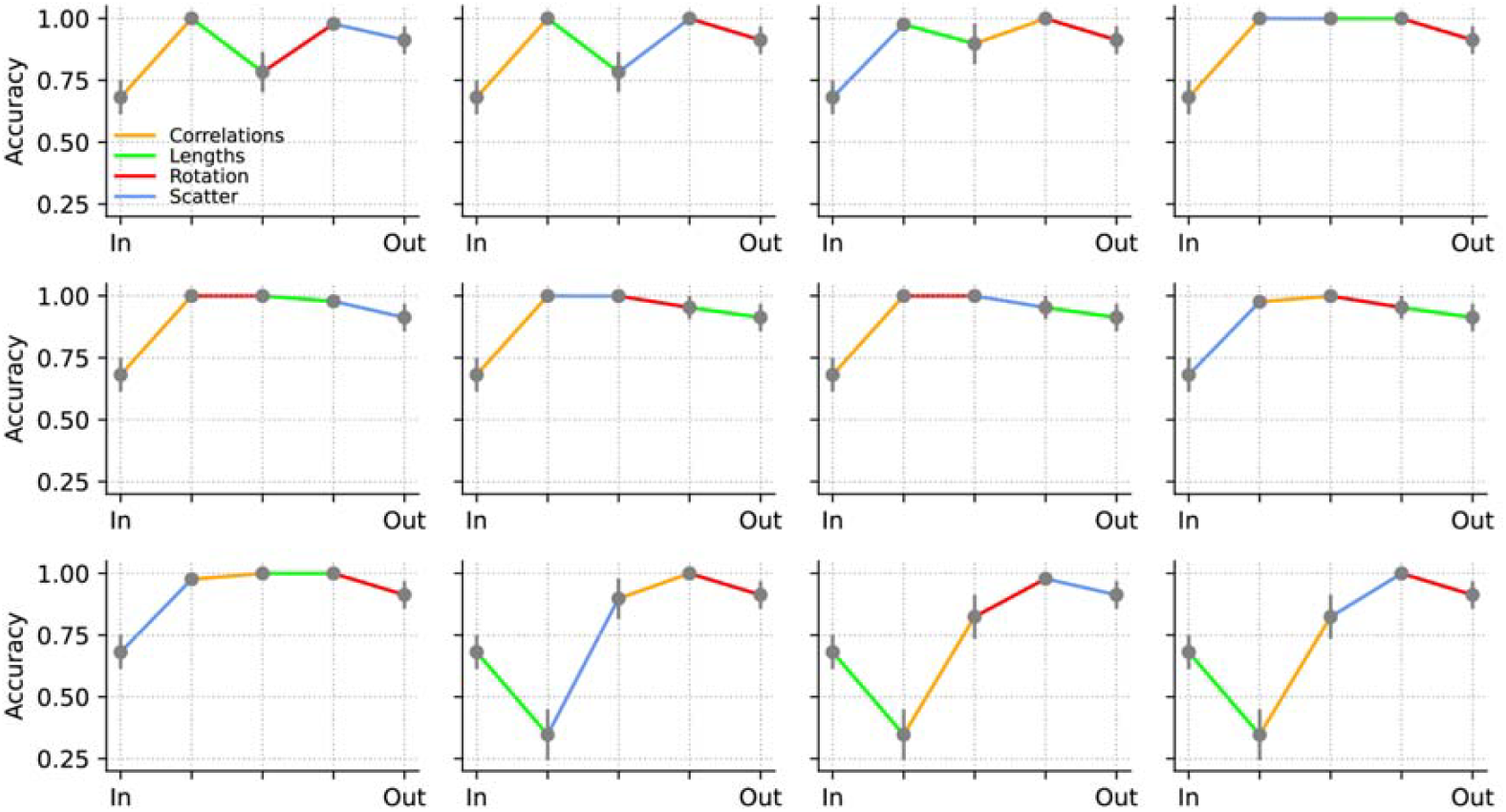
Relating decorrelation to accuracy by transforming input to output. The data in the first panel is shown in Figure 5 of the main text.

**Figure S5.3:**
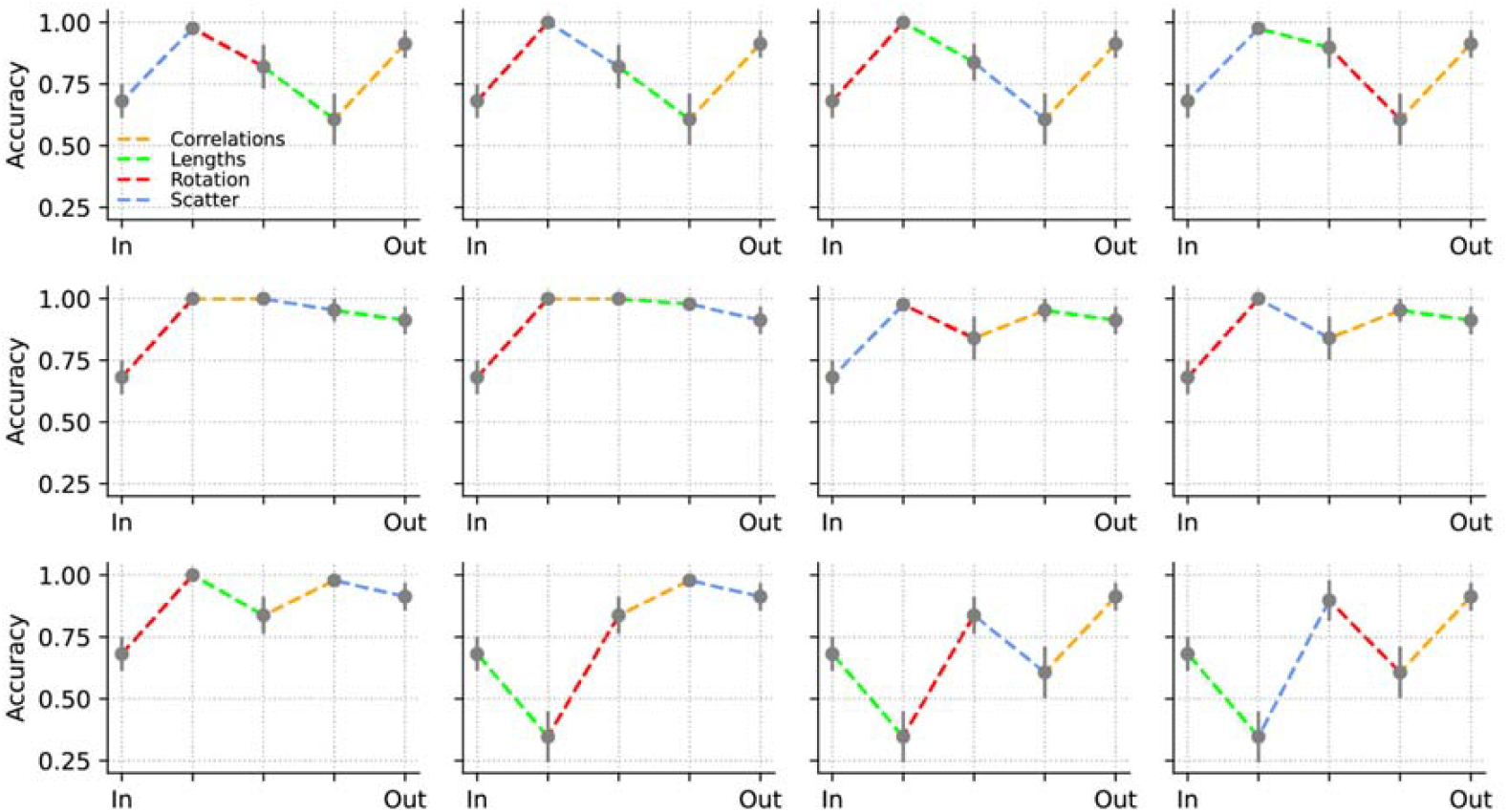
Relating decorrelation to accuracy by transforming output to input. The data in the first panel is shown in Figure 5 of the main text.

