## Supplementary figures (high res) for "Decorrelation by gain control in the mouse olfactory bulb"

### Figure S1.1

**A**

| Acids | Alcohols | Aldehydes | Aromatics | Esters | Ketones |
|---|---|---|---|---|---|
| Nonanoic acid | 1- Nonanol | 2-Phenylpropionaldehyde | 1,2 -Dimethoxybenzene | Ethyl Valerate | 2-Hydroxy acetophenone |
| Hexanoic acid | 1-Heptanol | Benzaldehyde | alpha-terpinene | Ethyl heptanoate | (+) -Fenchone |
| Isobutyric acid | 2-methyl -4-butanol | Valeraldehyde | 4-Allylanisole | Ethyl butyrate | 2-Nonanone |
| Valeric acid | Cyclohexanol | Octanal | R-(+) -Limonene | Ethyl acetate | 2-Heptanone |
| | | | 2-Methoxy-4-Methylphenol | cis -3-Hexenyl tiglate | Acetophenone |
| | | | Guaiacol | Propyl acetate | 4-Phenyl -2-Butanone |
| | | | Eugenol | Isoamyl acetate | S-(+) -Carvone |
| | | | Cineol | Methyl valerate | 2,4 -dimethyl acetophenone |
| | | | 1,4 -Cineol | Methyl salicylate | Menthone |
| | | | | Methyl butyrate | 2-Hexanone |
| | | | | Methyl benzoate | 4-methyl acetophenone |
| | | | | Geranyl acetate | |
| | | | | Ethyl Caproate | |
| | | | | Ethyl tiglate | |
| | | | | Benzyl acetate | |

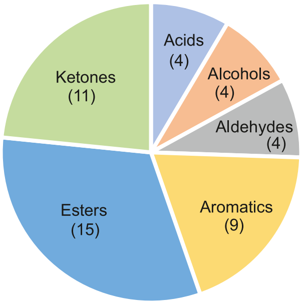


**B**

| Presentation order | | | |
|---|---|---|---|
| 0 | Nonanoic acid | 24 | Methyl salicylate |
| 1 | 2-Hydroxy acetophenone | 25 | 2-Methoxy-4-Methylphenol |
| 2 | 1-Nonanol | 26 | 2-Heptanone |
| 3 | 2-Phenylpropionaldehyde | 27 | cis-3-Hexenyl tiglate |
| 4 | 1,2-Dimethoxybenzene | 28 | Ethyl Caproate |
| 5 | Ethyl Valerate | 29 | Eugenol |
| 6 | 2-Nonanone | 30 | S-(+)-Carvone |
| 7 | 2-methyl-4-butanol | 31 | Methyl benzoate |
| 8 | Benzaldehyde | 32 | Octanal |
| 9 | Hexanoic acid | 33 | Valeric acid |
| 10 | Methyl valerate | 34 | Mineral Oil |
| 11 | Benzyl acetate | 35 | 2,4-dimethyl acetophenone |
| 12 | 1-Heptanol | 36 | Cineol |
| 13 | alpha-terpinene | 37 | Ethyl tiglate |
| 14 | Acetophenone | 38 | Isoamyl acetate |
| 15 | Valeraldehyde | 39 | Menthone |
| 16 | Geranyl acetate | 40 | 2-Hexanone |
| 17 | (+)-Fenchone | 41 | 4-Phenyl-2-Butanone |
| 18 | Ethyl heptanoate | 42 | Methyl butyrate |
| 19 | 4-Allylanisole | 43 | R-(+)-Limonene |
| 20 | Cyclohexanol | 44 | Ethyl butyrate |
| 21 | Propyl acetate | 45 | Ethyl acetate |
| 22 | 1,4-Cineol | 46 | 4-methyl acetophenone |
| 23 | Guaiacol | 47 | Isobutyric acid |

**C**

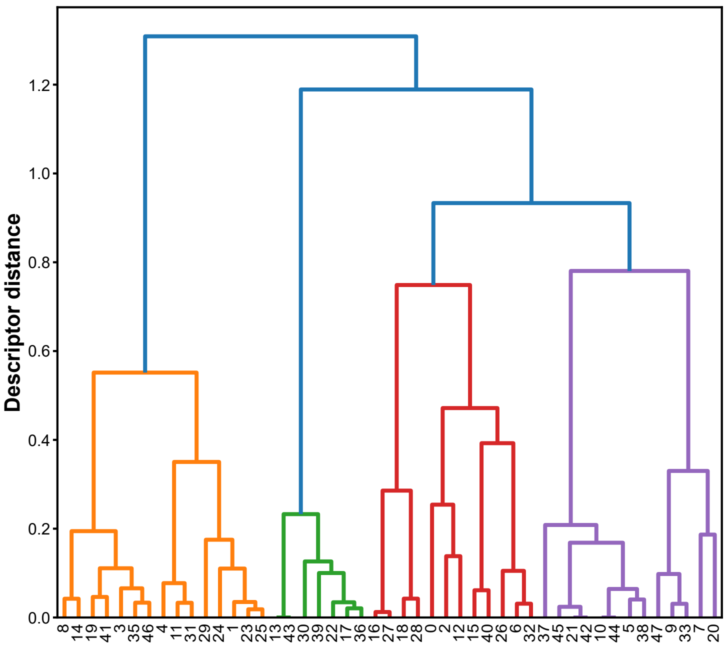


**Figure S1.1: Experimental odour panel. (A)** List of monomolecular odours presented during experiments sorted by chemical class. **(B)** Sequences of odours as presented during experiment. **(C)** Molecular descriptor-based clustering of odour panel. Hierarchical clustering dendrogram based on 9 molecular descriptors (correlation distance, average linkage). Colors indicate chemical class. Numbers show odour IDs.

### Figure S1.2

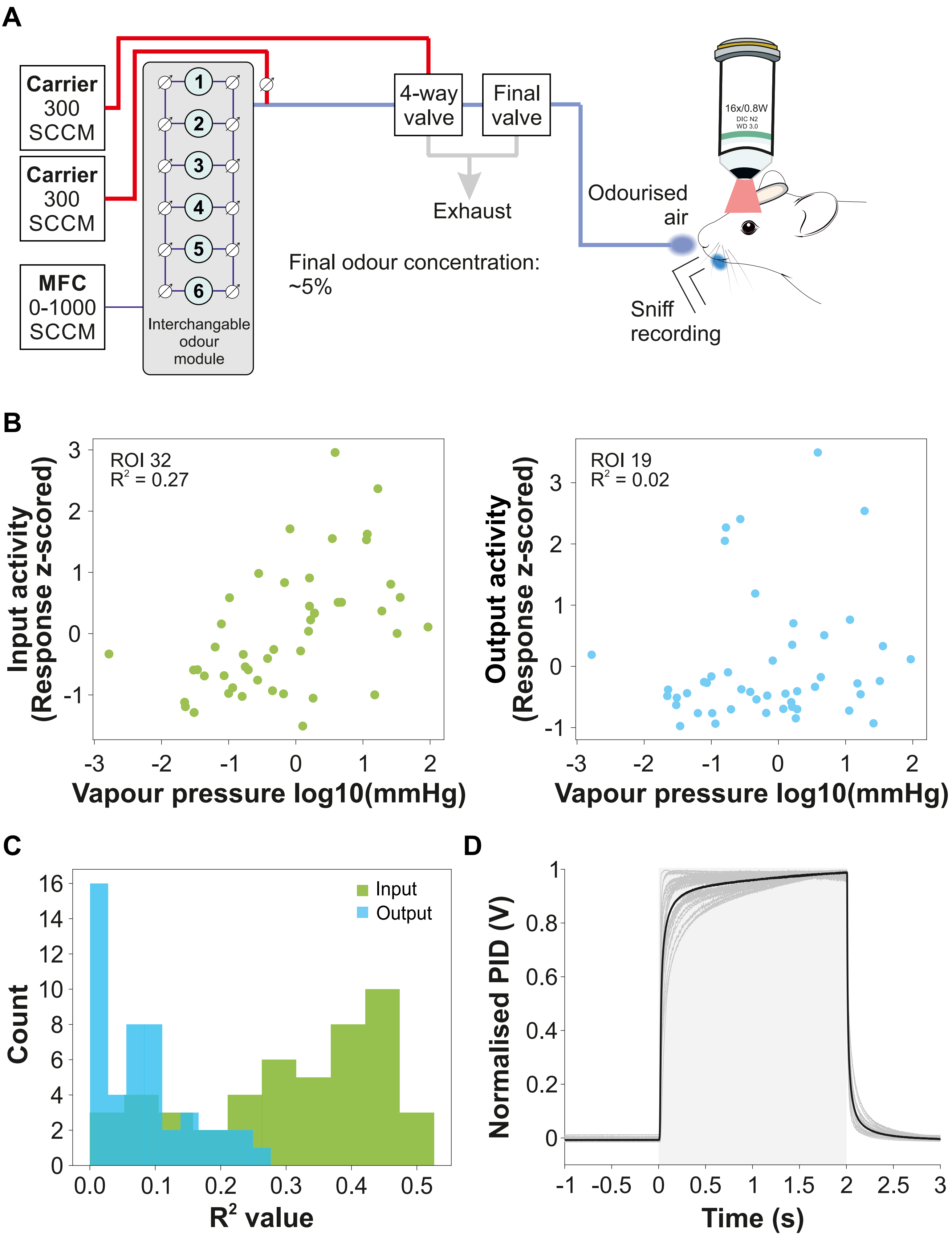


**Figure S1.2: Schematic of odour presentation and stimulus characteristics. (A)** Schematic of the custom odour delivery system. **(B)** Z-scored odour response of an example input (left) and output glomerulus (right) plotted as a function of odour vapour pressure shown in a log10 scale. **(C)** Histogram of R-squared values for all glomeruli computed by linearly regressing each glomerular z-scored response as a function of vapour pressure. **(D)** Overlay of temporal profiles of 21 odours as recorded using photoionisation detector (PID). Shown are recordings of all 21 monomolecular odours that elicited a voltage signal in the PID (grey traces) and the average signal of these odours (black trace). Odour presentation period from 0-2 seconds (grey shaded area), traces are normalised between 0 and 1.

### Figure S2.1

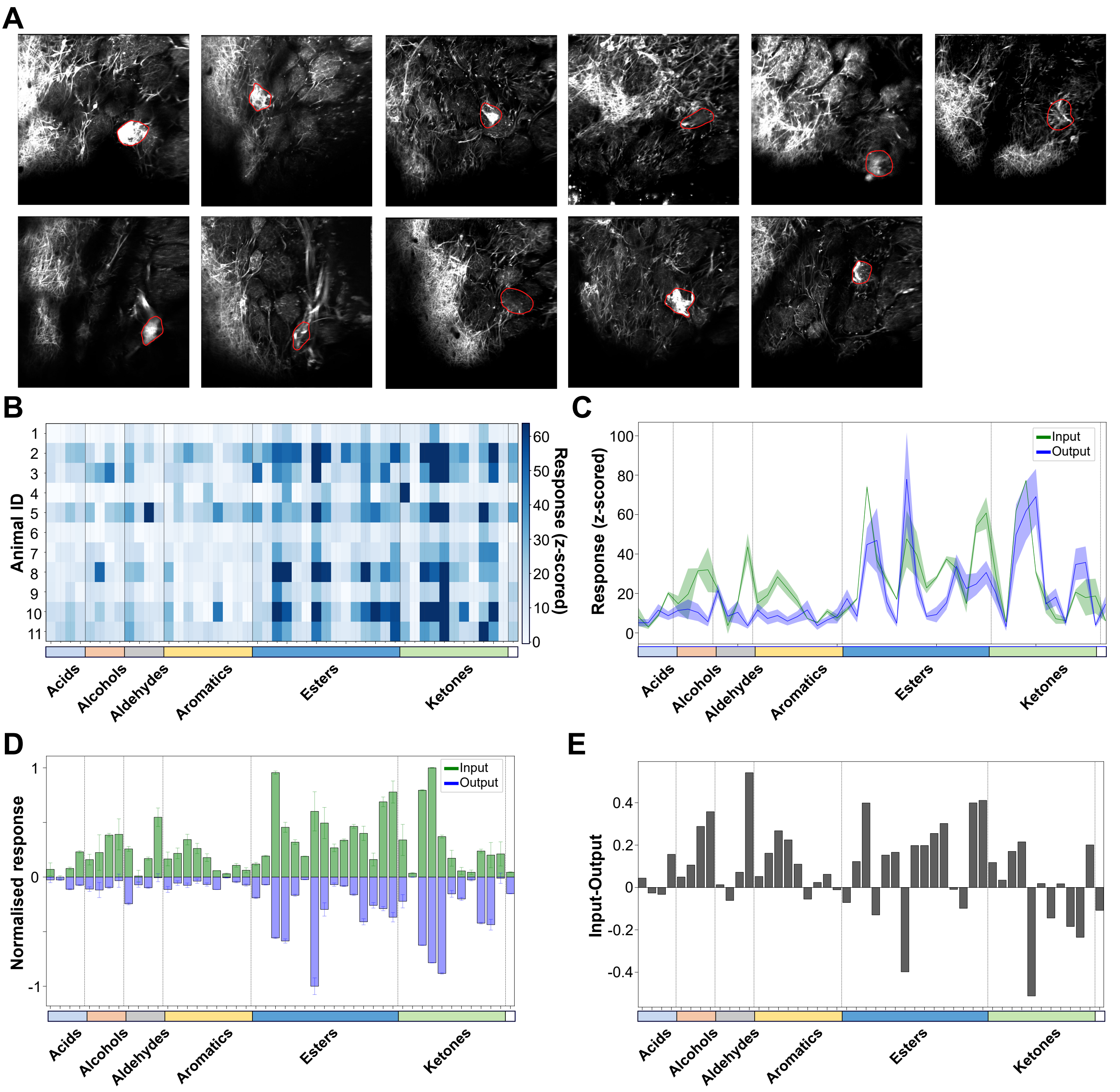


**Figure S2.1: Input–output transformation of M72 glomerular odour responses. (A)** GCaMP6f fluorescence from output neuron dendrites recorded in the glomerular layer of Tbet-cre:M72-ChR2-YFP:Ai95 mice. The M72 glomerulus is identified by YFP-labelled sensory axons (red contour; M72-YFP; n = 11 animals). **(B)** Heatmap of individual M72 output responses (z-scored and integrated) to all odours sorted by chemical class**. (C)** Average z-scored responses to all odours as measured from the M72 glomerulus (mean +/- SEM, input: green, output: blue). **(D)** Normalised average responses to all odours presented as mean +/- SD (input: green, plotted as positive; output: blue, plotted as negative). **(E)** M72 response profile plotted as difference between input and output signal. Input: n = 2 animals, output: 11 animals.

### Figure S3.1

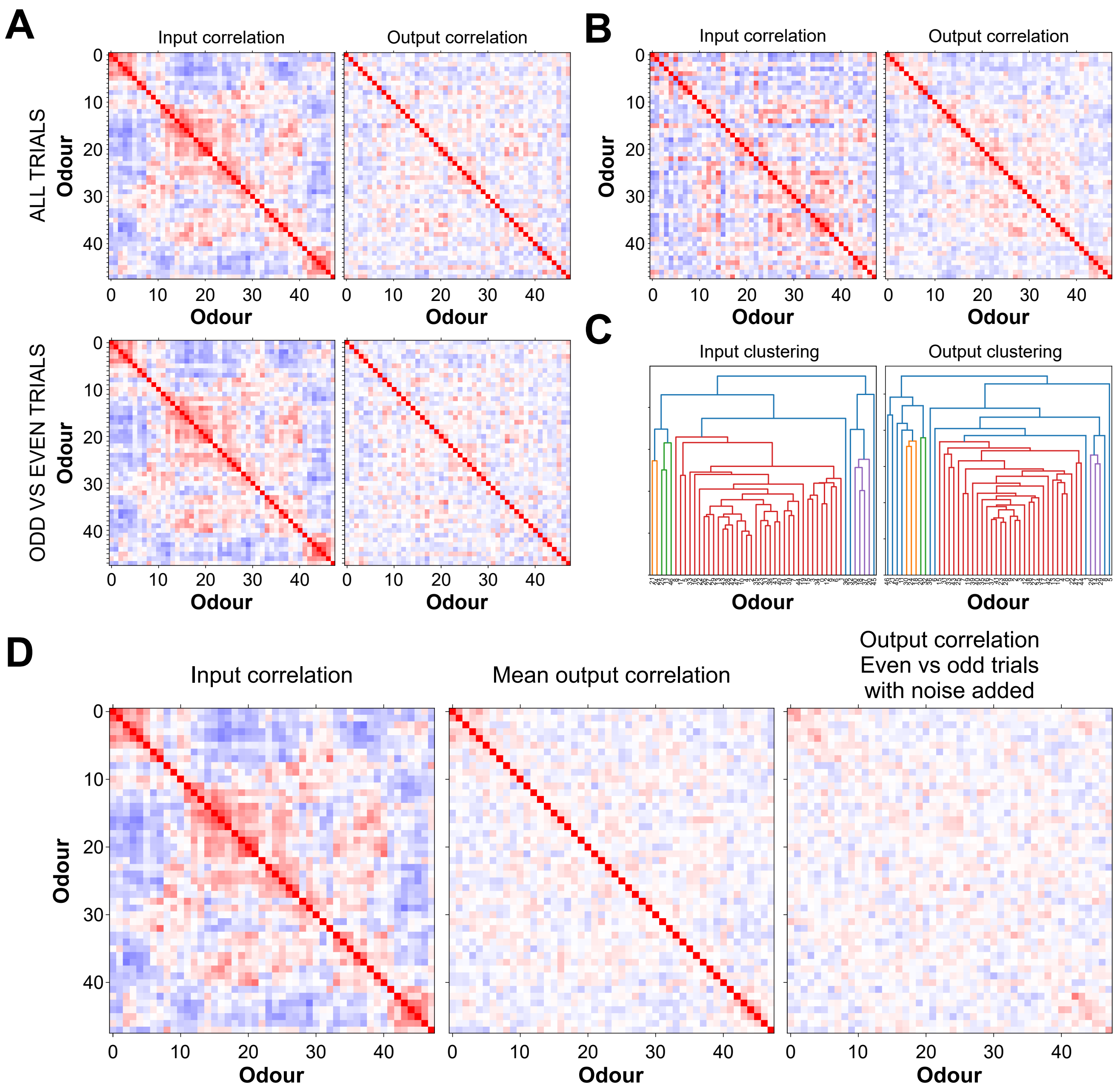


**Figure S3.1: Decorrelation is consistent across trials and is not based on random noise. (A)** Correlation matrices for input and output data computed by using all trials (top row) and by correlating odd against even trials. **(B)** Correlation matrices using odour sequence determined by hierarchical clustering based on the Euclidean distance of the output data. **(C)** Dendrograms resulting from hierarchical clustering of input (left) and output data (right). **(D)** The addition of random noise (2 s.d.) to the raw data before computing the correlation matrix of the output data reduces the cross correlation to approximately zero on the diagonal, proving that the decorrelation is not down to random noise.

### Figure S3.2

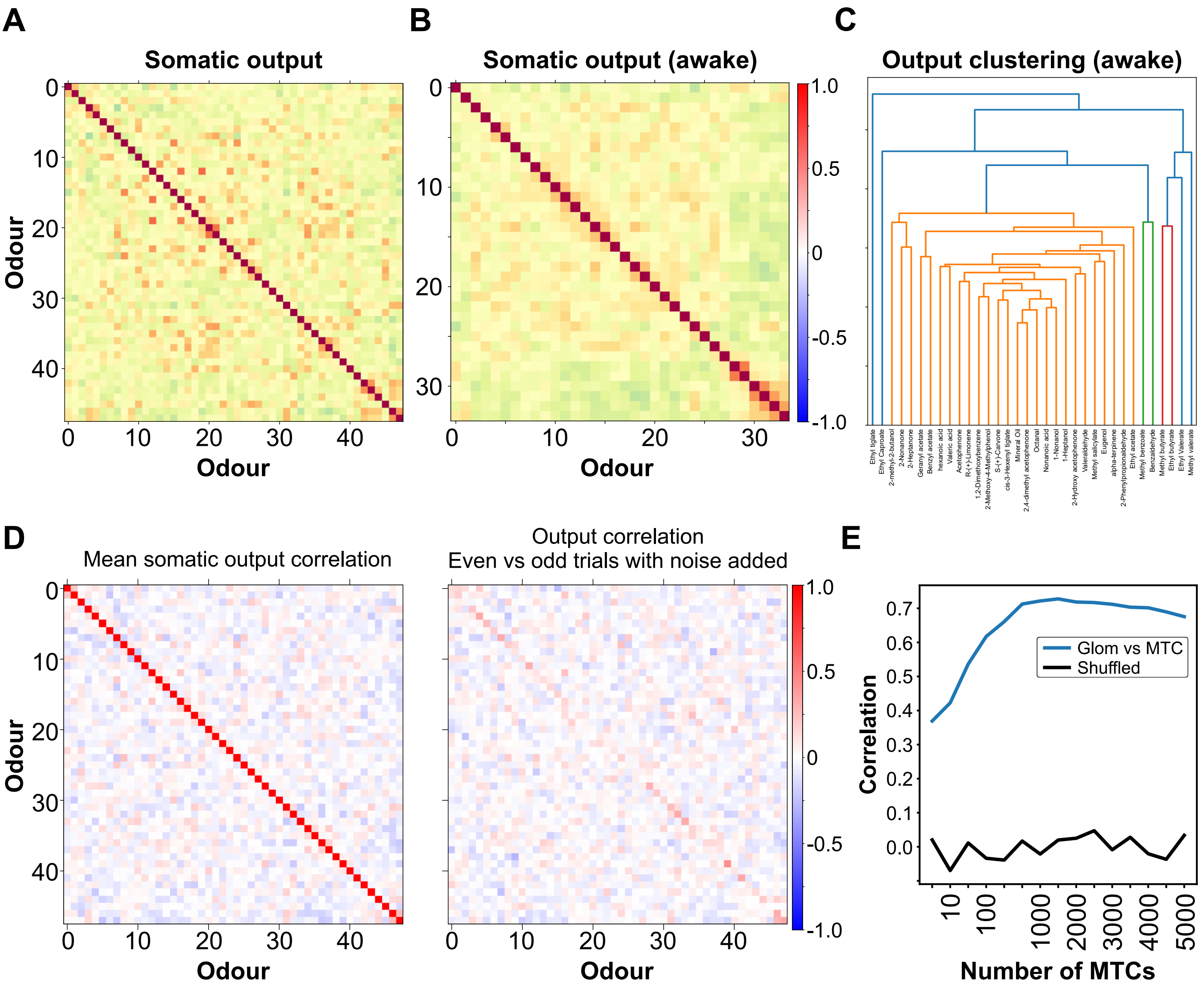


**Figure S3.2: Pattern correlation of somatic odour responses. (A)** Odour correlation maps of somatic ROIs. The sequence of odours was determined using hierarchical clustering based on the Euclidean distance of the input data. To equalise the number of ROIs for both datasets, 167 somatic ROIs that showed the highest variance across odour responses were selected. **(B)** Odour correlation (34 odours) for somatic ROIs (n = 167 MTCs) recorded in awake mice, rendering a comparable correlation structure to the anaesthetised state. **(C)** Hierarchical clustering used to define the sequence of odours shown in (B). **(D)** The addition of random noise (2 s.d.) to the correlation matrix of the output data reduces the cross correlation to approximately zero on the diagonal, proving that the decorrelation is not down to random noise. **(E)** Correlation between glomerular and somatic correlation matrices as a function of the number of M/T cells that were used. The correlation was calculated using the off-diagonal values of correlation matrices (blue: original order, black: order of correlation shuffled row wise).

### Figure S3.3

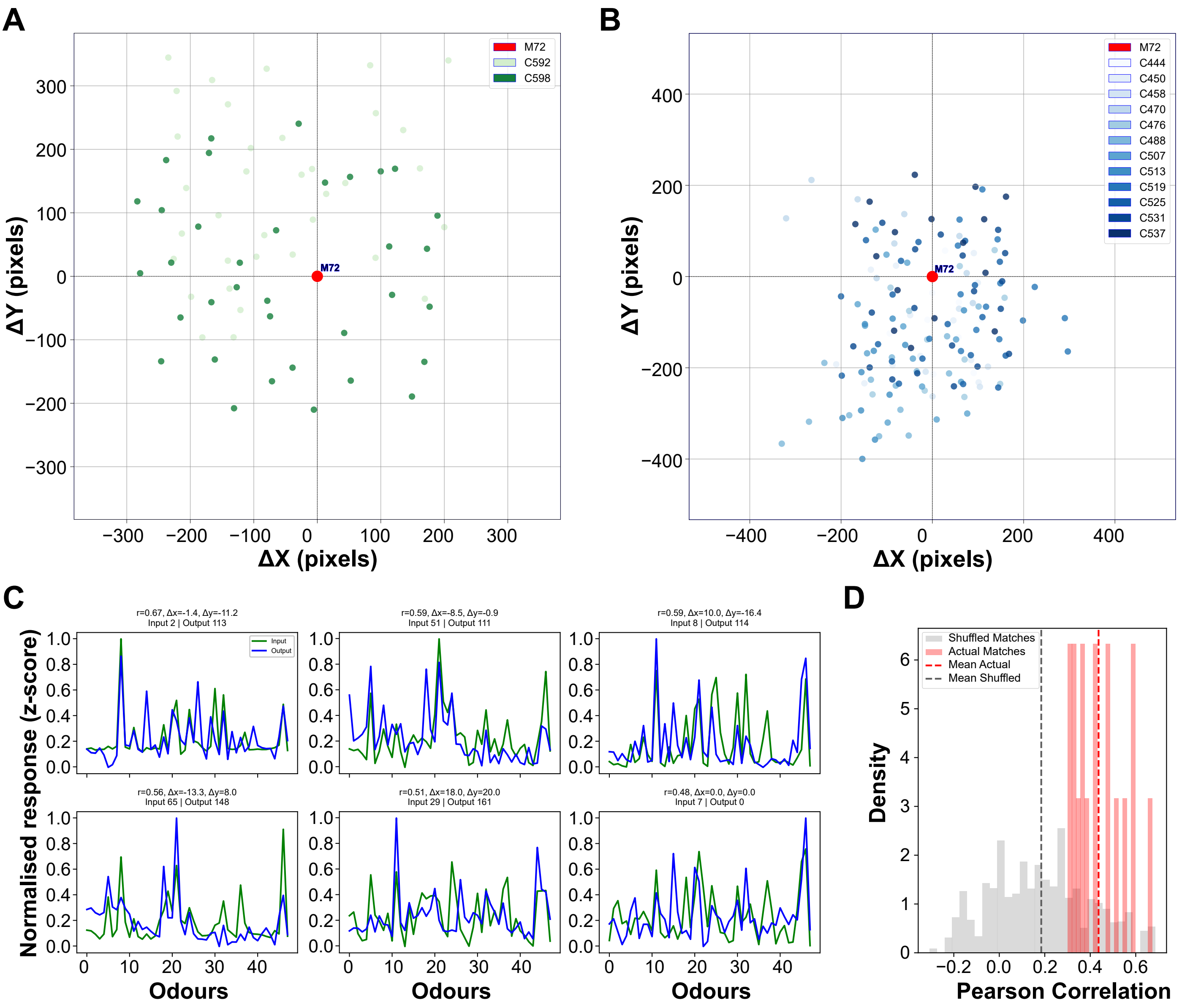


**Figure S3.3: Spatial and functional matching of glomeruli centered around the M72 glomerulus. (A)** Spatial distribution of glomerular input ROIs from two animals, aligned to the centroid of the M72 glomerulus (red). Colour intensity reflects different animals. **(B)** Same as in (A), but showing the distribution of output ROIs from twelve animals. **(C)** Odour response profiles of nine representative input–output ROI matched pairs with high correlation, sorted by descending Pearson correlation (r). Input and output responses are shown in green and blue, respectively. **(D)** Distribution of Pearson correlation coefficients for functionally matched ROI pairs with $r > 0.3$ (red) and shuffled controls (grey). Matched pairs showed significantly higher correlation than shuffled controls (Kolmogorov-Smirnov test, $p = 1.73e\text{-}09$; KS statistic = 0.721), suggesting consistent matching across input and output representations centered on M72.

### Figure S3.4

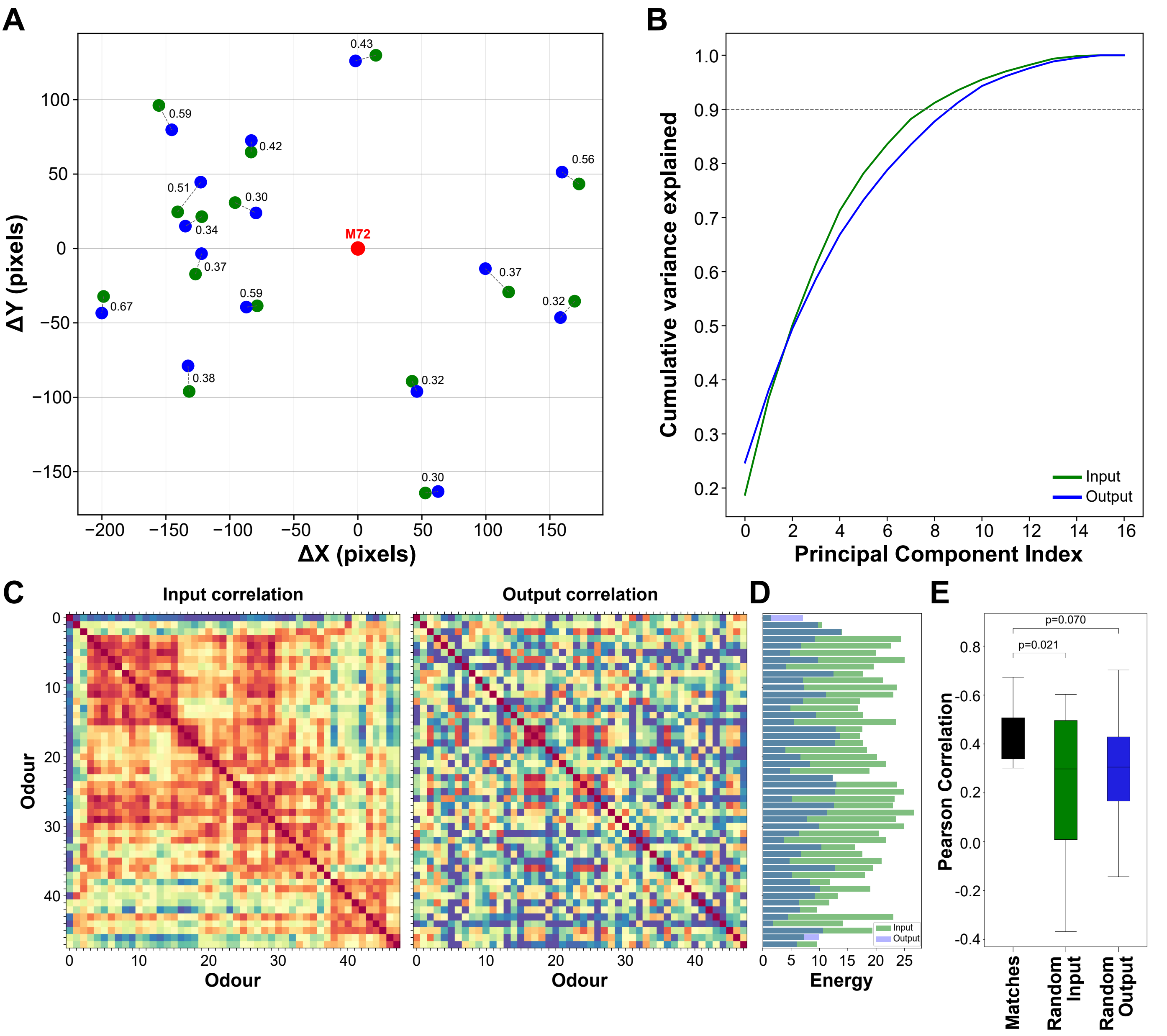


**Figure S3.4: Odour representations of matched glomeruli become decorrelated from input to output. (A)** Spatial distribution of input (green) and output (blue) ROIs matched by both spatial proximity and functional similarity (Pearson correlation r > 0.3), aligned to the M72 glomerulus (red) as a common reference point. Correlation values for each matched pair are indicated. **(B)** Cumulative variance explained by principal component analysis (PCA) of matched glomerular odour responses. Input responses (green) reached 90% variance with 9 components compared to output responses (blue, PC 10), indicating larger dimensionality in output representations for the subset of matched glomeruli. **(C)** Odour correlation matrices for matched input (left) and output (right) glomeruli (n = 17). Odour order was fixed across both matrices using hierarchical clustering based on the input data to facilitate direct comparison. The output matrix exhibits less structure and lower correlation, consistent with decorrelation. **(D)** Energy of odour correlations (defined as the squared sum of pairwise correlations minus 1) for each odour, comparing input (green) and output (blue). Most odours show reduced correlation energy in the output, further supporting a broad decorrelation of glomerular representations. **(E)** Correlation values between spatially matched glomeruli are significantly higher than correlations with randomly paired input or output ROIs (n = 17 ROIs).

### Figure S4.1

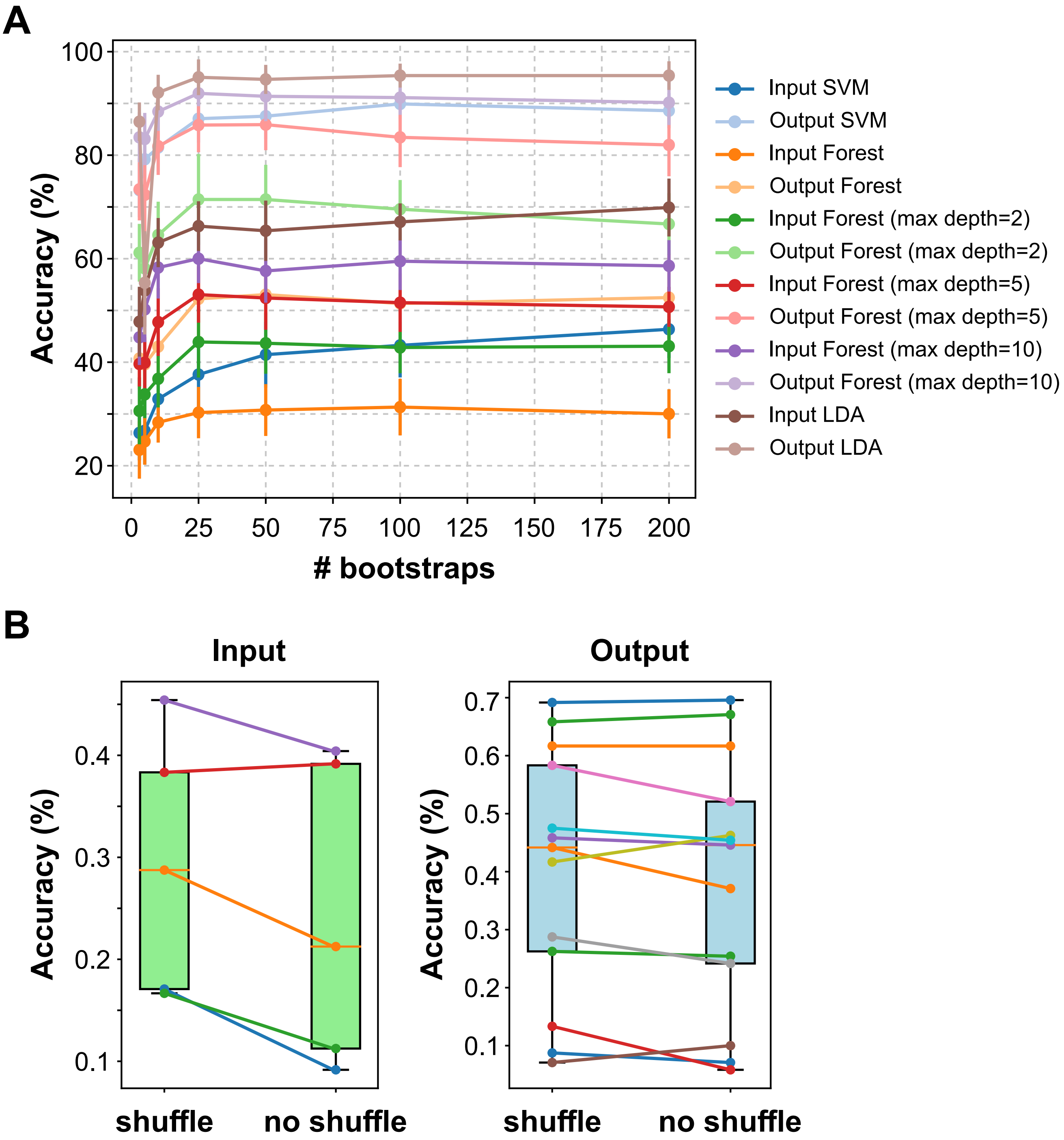


**Figure S4.1: Effect of bootstrapping and label shuffling on classifier accuracy. (A)** Comparison of different classifiers (SVM, Random Forest and LDA) on odour prediction accuracy as a function of the number of bootstrapped odour repetition trials. Accuracy plateaus for most classifiers at around 25 trials. **(B)** Shuffling trial order and thereby omitting noise correlations when using 3-5 experimentally acquired trials has no significant effect on prediction accuracy (input shuffle: 0.29 ± 0.11, input no shuffle: 0.24 ± 0.13, mean ± s.d., p = 0.54; output shuffle: 0.42 ± 0.21, output no shuffle: 0.39 ± 0.21, mean ± s.d., p = 0.78).

### Figure S5.1

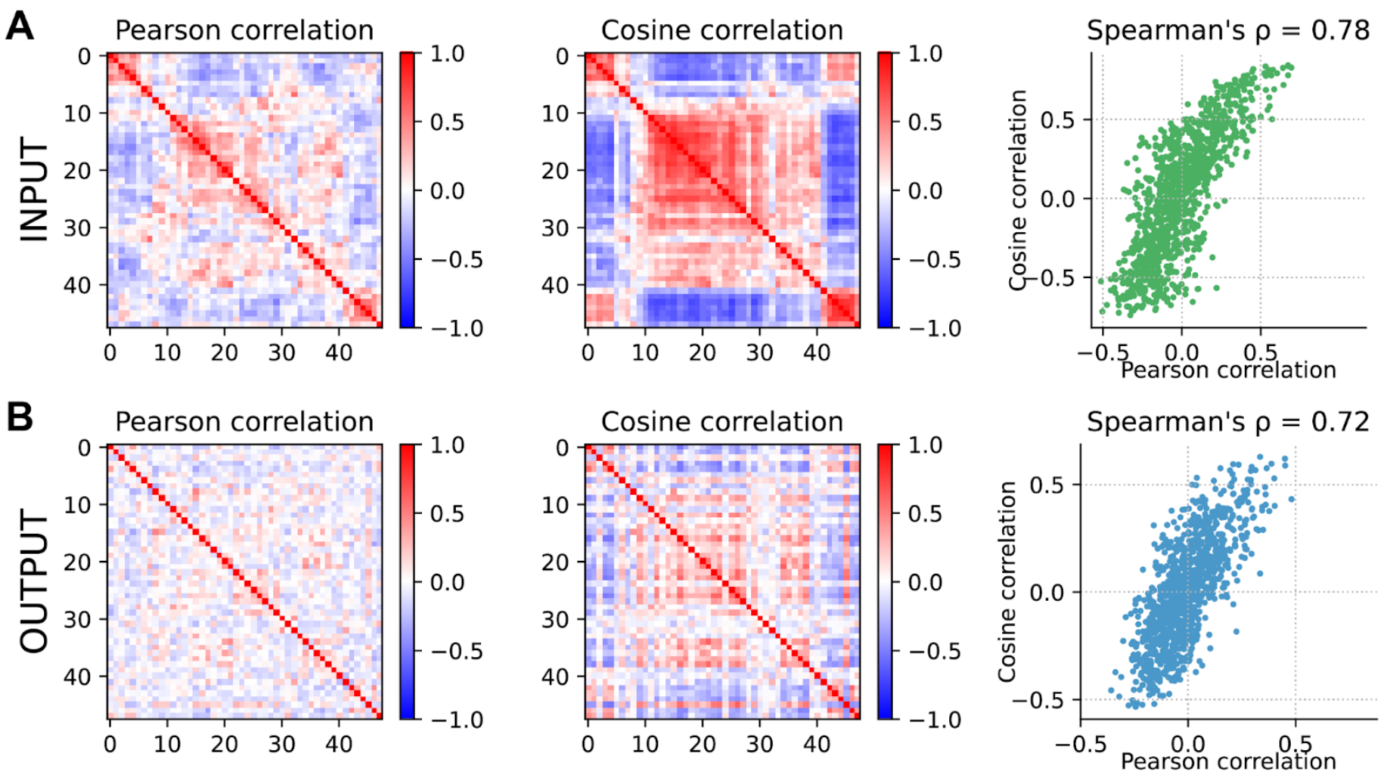


**Figure S5.1: Pearson correlation and cosine correlation are correlated. (A)** Pearson correlations (left) and cosine correlations (middle) of trial-averaged odour responses of olfactory bulb inputs. Ticks indicate odours, ordered as in the main text. Pearson correlation plots are as in the Main Text; right: Pearson correlation vs. cosine correlation for all pairs of odours. **(B)** As in (A) but for olfactory bulb outputs.

### Figure S5.2

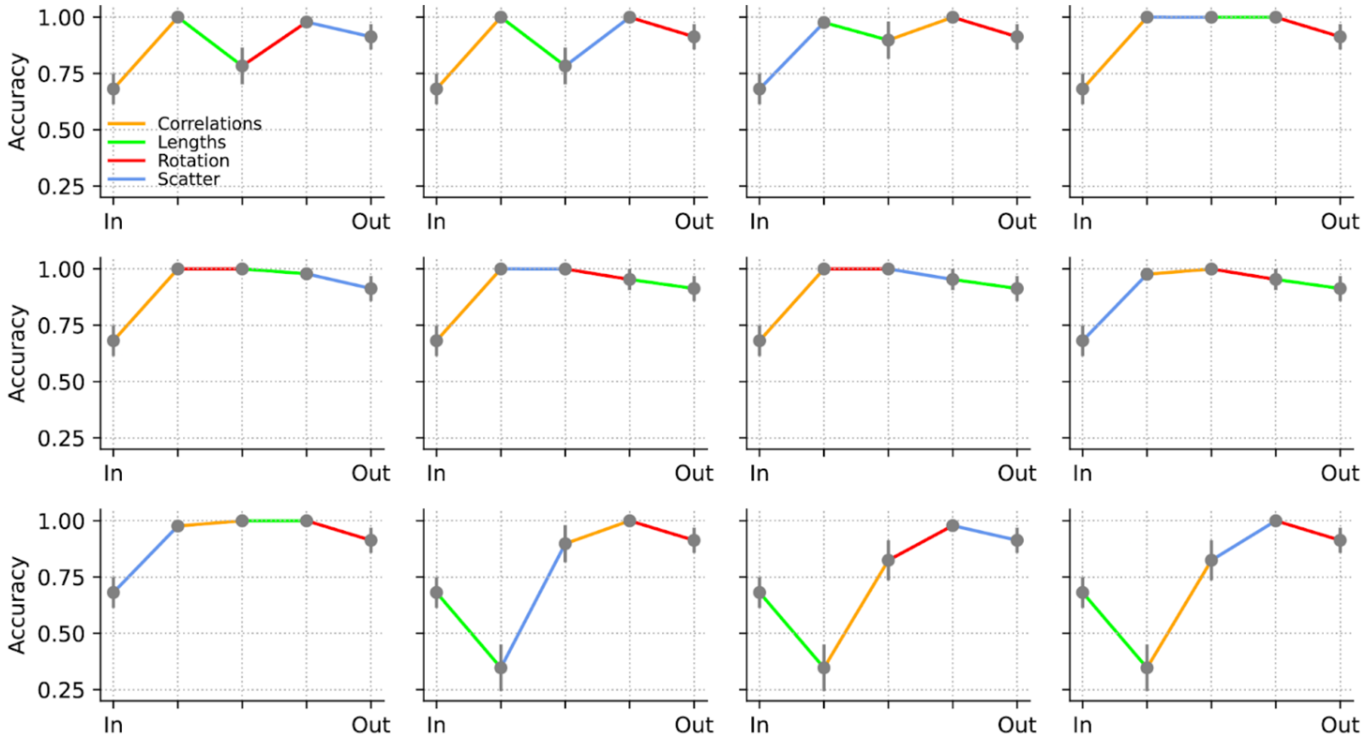


**Figure S5.2: Relating decorrelation to accuracy by transforming input to output.** The data in the first panel is shown in Figure 5 of the main text.

### Figure S5.3

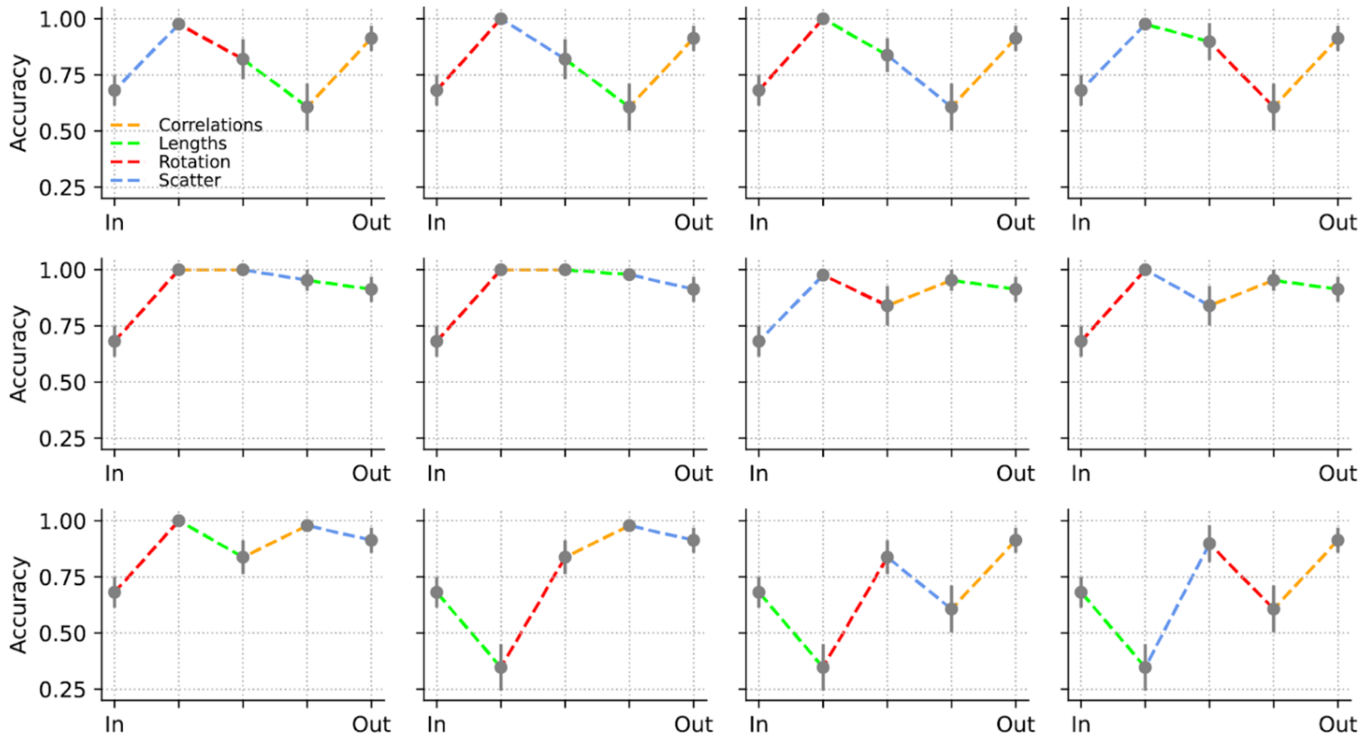


**Figure S5.3: Relating decorrelation to accuracy by transforming output to input.** The data in the first panel is shown in Figure 5 of the main text.
